# DRAM for distilling microbial metabolism to automate the curation of microbiome function

**DOI:** 10.1101/2020.06.29.177501

**Authors:** Michael Shaffer, Mikayla A. Borton, Bridget B. McGivern, Ahmed A. Zayed, Sabina L. La Rosa, Lindsey M. Solden, Pengfei Liu, Adrienne B. Narrowe, Josué Rodríguez-Ramos, Benjamin Bolduc, M. Consuelo Gazitua, Rebecca A. Daly, Garrett J. Smith, Dean R. Vik, Phil B. Pope, Matthew B. Sullivan, Simon Roux, Kelly C. Wrighton

## Abstract

Microbial and viral communities transform the chemistry of Earth’s ecosystems, yet the specific reactions catalyzed by these biological engines are hard to decode due to the absence of a scalable, metabolically resolved, annotation software. Here, we present DRAM (<u>D</u>istilled and <u>R</u>efined <u>A</u>nnotation of <u>M</u>etabolism), a framework to translate the deluge of microbiome-based genomic information into a catalog of microbial traits. To demonstrate the applicability of DRAM across metabolically diverse genomes, we evaluated DRAM performance on a defined, *in silico* soil community and previously published human gut metagenomes. We show that DRAM accurately assigned microbial contributions to geochemical cycles, and automated the partitioning of gut microbial carbohydrate metabolism at substrate levels. DRAM-v, the viral mode of DRAM, established rules to identify virally-encoded auxiliary metabolic genes (AMGs), resulting in the metabolic categorization of thousands of putative AMGs from soils and guts. Together DRAM and DRAM-v provide critical metabolic profiling capabilities that decipher mechanisms underpinning microbiome function.

## INTRODUCTION

DNA sequencing advances have offered new opportunities for cultivation-independent assessment of microbial community membership and function. Initially, single gene approaches established taxonomic profiling capabilities, providing innumerable intellectual leaps in microbial composition across biomes (1, 2). Recently, the field has expanded from gene-based methods towards metagenome-assembled-genome (MAG) studies, which offer population level inferences of microbial functional underpinnings (3–5). Across ecosystems, these MAGs illuminated new biological feedbacks to climate-induced changes (6–8), revolutionized personalized microbiota-based therapeutics for human health (9, 10), and dramatically expanded the tree of life (11–13). Metagenomic advances have also transformed our ability to study viruses, and since they lack a universal barcode gene, viral MAG (vMAG) enabled studies are required for even viral taxonomic surveys (14, 15).

At this point, there are hundreds of thousands of MAGs and vMAGs available from the human gut and other diverse environments (7, 14–23). This inundation of data required development of scalable, genome-based taxonomic approaches, which are now largely in place for both microbes (24, 25) and viruses (26, 27). However, there is a growing consensus that for any of these habitats the taxonomic composition of the microbiome alone is not a good predictor of ecosystem functions, properties which are often better predicted from microbial and viral traits (28, 29). Therefore, there is an absolute need to develop gene annotation software that can simultaneously highly resolve trait prediction from vast amounts of genomic content.

While there are several tools for annotating genes from microbial genomes (30–33), a single tool has yet to translate current knowledge of microbial metabolism into a format that can be applied across thousands of genomes. Most online annotators are only useful for a handful of genomes or for profiling genes using a single database (34–36). Other recently developed tools have advanced to annotate thousands of genomes with multiple databases, which expands the biological information queried (30–32). However, biological interpretation is still burdened by challenges in data synthesis and visualization, thereby preventing efficient metabolic profiling of microbial traits with known ecosystem relevance. In addition, viruses can encode Auxiliary Metabolic Genes (AMGs) that directly reprogram key microbial metabolisms like photosynthesis, carbon metabolism, and nitrogen and sulfur cycling (37, 38), but identifying and insuring these AMGs are not ‘contaminating’ microbial DNA (39) remains a painfully manual process.

Here we present a new tool, DRAM (Distilled and Refined Annotation of Metabolism), and the companion tool DRAM-v for viruses, and apply these tools to existing, assembled metagenomic datasets to demonstrate the expanded utility over past approaches. DRAM was designed to profile microbial (meta)genomes for metabolisms known to impact ecosystem function across biomes and is highly customizable to user annotations. DRAM-v leverages DRAM’s functional profiling capabilities, and adds a ruleset for defining and annotating AMGs in viral genomes. Together DRAM and DRAM-v decode the metabolic functional potential harbored in microbiomes.

## MATERIAL AND METHODS

### DRAM annotation overview

The DRAM workflow overview is detailed in **Figure 1**. DRAM does not use unassembled reads, but instead uses assembly-derived FASTA files input by the user. Input files may come from unbinned data (metagenome contig or scaffold files) or genome-resolved data from one or many organisms (isolate genomes, single-amplified genome (SAGs), MAGs). First each file is filtered to remove short contigs (by default contigs <2500bp, but this can be user defined). Then Prodigal (40) is used to detect open reading frames (ORFs) and subsequently predict their amino acid sequences, supporting all genetic codes on defined on NCBI (**Figure 1, Supplementary Figure 1**). Specifically, we use the anonymous/metagenome mode of Prodigal (40), which is recommended for metagenome assembled contigs and scaffolds. By default, first Prodigal (40) tests genetic code 11, then uses other genetic codes to resolve short genes, or notifies user that no code resolves gene length.

**Figure 1:**
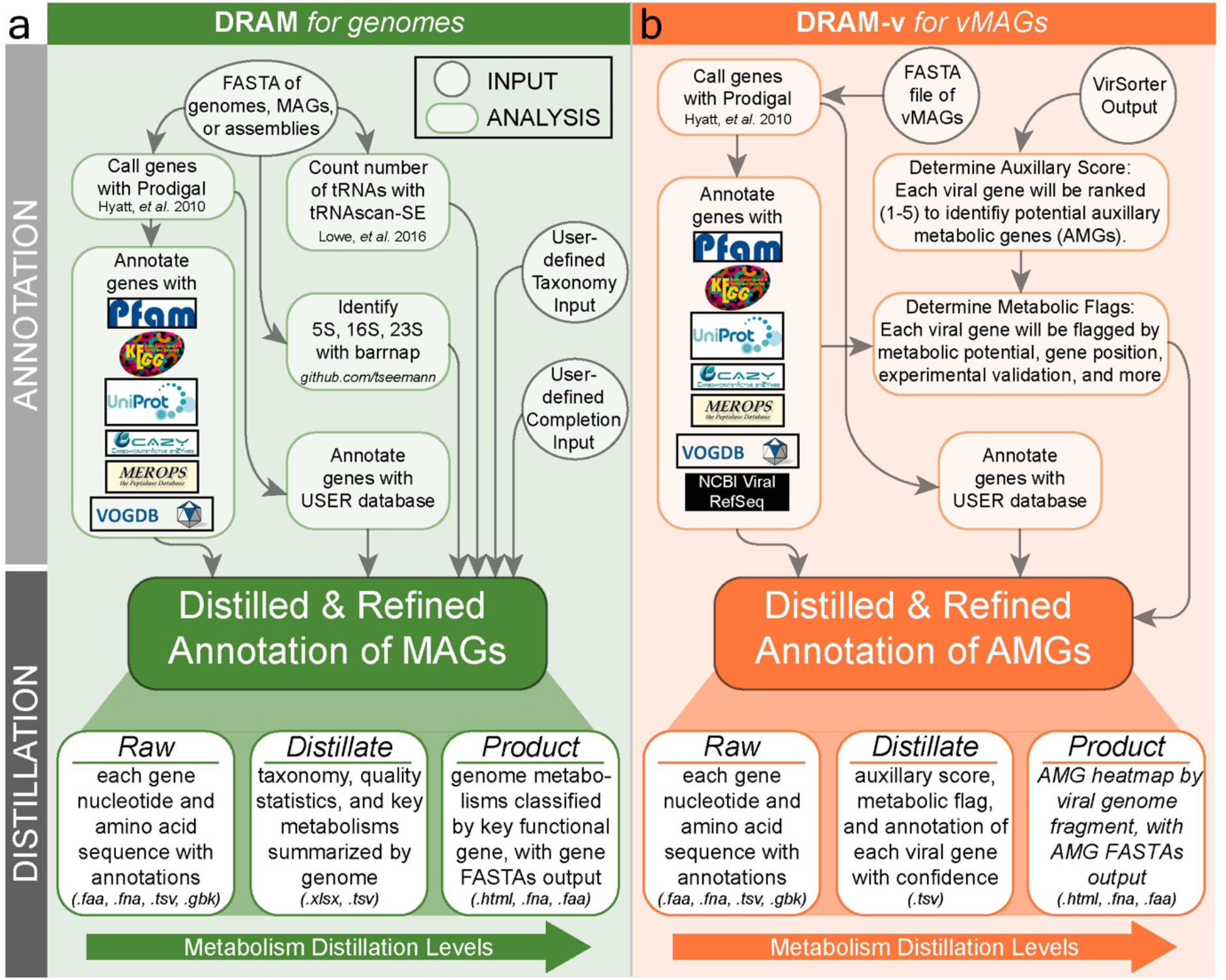
Conceptual overview and workflow of the assembly-based software, DRAM (Distilled and Refined Annotation of Metabolism). DRAM (green, **a**) profiles microbial metabolism from genomic sequences, while DRAM-v profiles the Auxiliary Metabolic Genes (AMGs) (orange, **b**) in vMAGs. DRAM’s input data files are denoted by circles in grey, while analysis and output files are denoted by rectangles in green for MAGs or orange for AMGs. DRAM’s outputs (from the *Raw, Distillate*, and *Product*) provide three levels of annotation density and metabolic parsing. More details on the output files and specific operation can be found in the **Supplementary Text** or at https://github.com/shafferm/DRAM/wiki. User defined taxonomy (e.g. GTDB-Tk (24)) and completion estimates (e.g. CheckM (51)) for MAGs and isolate genomes can be input into DRAM.

Next, DRAM searches all amino acid sequences against multiple databases and provides all database hits in a single output file called the *Raw* output (**Supplementary File 1, Supplementary Figure 1**). Specifically, ORF predicted amino acid sequences are searched against KEGG (41), Uniref90 (42), and MEROPS (43) using MMseqs2 (44), with the best hits (defined by bitscore, default minimum threshold of 60) reported for each database in the *Raw* output. Note, the use of the Uniref90 (42) database is not default due to the increased memory requirements which can be prohibitive to many users, thus a user should specify the --use_uniref flag to search amino acid sequences against Uniref90 (42). If there is no hit for a given gene in a given database above the minimum bit score threshold, no annotation is reported for the given gene (unannotated) and database in the *Raw* output. Reciprocal best hits (RBHs) are defined by searches where the database sequence that is the top hit from a forward search of the input gene has a bit score greater than 60 (by default) and is the top hit from the reverse search of the database hit against the all genes from the input FASTA file with a bit score greater than 350 (by default) (3, 45). DRAM also uses MMSeqs2 (44) to perform HMM profile searches of the Pfam database (46), while HHMER3 (47) is used for HMM profile searches of dbCAN (48) and VOGDB (http://vogdb.org/). For these HMM searches of Pfam, dbCAN, and VOGDB, a hit is recorded if the coverage length is greater than 35% of the model and the e-value is less than 10^−15^ (48). If the user does not have access to the KEGG database, DRAM automatically searches the KOfam (49) database with HMMER in order to assign KOs, using gene specific e-value and percent coverage cutoffs provided here ftp://ftp.genome.jp/pub/db/kofam/ko_list.gz (49). Users should note that using KOfam (49) rather than KEGG genes (41), may result in less annotation recovery, thereby resulting in some false negatives in the DRAM *Product* (described below). After ORF annotation, tRNAs are detected using tRNAscan-SE (50) and rRNAs are detected using barrnap (https://github.com/tseemann/barrnap).

When gene annotation is complete, the results are merged to a single tab-delimited annotation table that includes the best hit from each database for user comparison. (**Supplementary File 1, Supplementary Figure 1**). For each gene annotated, DRAM provides a single, summary rank (A-E), which represents the confidence of the annotation (**Supplementary Figure 1**). The highest rank includes reciprocal best hits (RBH) with a bit score >350, against KEGG (41) genes (A rank) (41), followed by reciprocal best hits to Uniref90 (42) with a bit score >350 (B rank), hits to KEGG (41) genes (41) with a bit score >60 (C rank), and UniRef90 (42) with a bit score greater than 60 (C rank) (45). The next rank represents proteins that only had Pfam (46), dbCAN (48), or MEROPS (43) matches (D rank), but hits to KEGG (41) or UniRef90 (42) were below 60 bit score. The lowest rank (E) represents proteins that had no significant hits to any DRAM database including KEGG (41), Uniref90 (42), dbCAN (48), Pfam (46), MEROPS (43), or only had significant hits to VOGDB. **Supplementary Figure 1** provides a schematic summarizing this annotation system. If one or more of the databases used for determining annotation ranks (KEGG, Uniref90, Pfam) is not used during DRAM annotation, all genes are considered to not have any hits against the unused database(s) and the respective annotation rank (e.g. B in the case of UniRef90) would be absent depending on which database was not used. In summary, the *Raw* output of DRAM provides for each gene in the dataset a summary rank (A-E), as well as the hits across up to 6 databases including KEGG, Uniref90, Pfam, CAZY, MEROPS, and VOGDB, allowing users to easily compare annotation content provided by different sources.

Beyond annotation, DRAM is intended to be a data compiler. Users can provide output files from GTDB-tk (24) and checkM (51) (or other user defined taxonomy and completion estimates), which are input into DRAM to provide taxonomy and genome quality information of the MAGs, respectively. For downstream analyses, DRAM provides a FASTA file of all entries from all input files, a GFF3-formatted file containing all annotation information, FASTA files of nucleotide and amino acid sequences of all genes, and text files with the count and position of the detected tRNAs and rRNAs (**Supplementary Figure 1**). Finally, a folder containing one GenBank formatted file for each input FASTA is created.

DRAM *Raw* annotations are distilled to create genome statistics and metabolism summary files, which are found in the *Distillate* output (**Supplementary File 2**). The genome statistics file provides most genome quality information required for MIMAG (25) reporting, including GTDB-tk (24) and checkM (51) information, if provided by the user. The summarized metabolism table contains the number of genes with specific metabolic function identifiers (KO, CAZY ID etc.) for each genome, with information distilled from multiple sources, including custom-defined metabolism modules (see https://raw.githubusercontent.com/shafferm/DRAM/master/data/genome_summary_form.tsv). For ease of metabolic interpretation, in the *Distillate*, many of the genes annotated in the *Raw* that can be assigned to pathways are output to multiple sheets assigned by functional category and organized by pathway (e.g. energy, carbon utilization, transporters) (**Figure 2ab**). Thus, the *Distillate* provides users with a pathway-centric organization of genes annotated in the *Raw*, while also summarizing the genome quality statistics.

**Figure 2:**
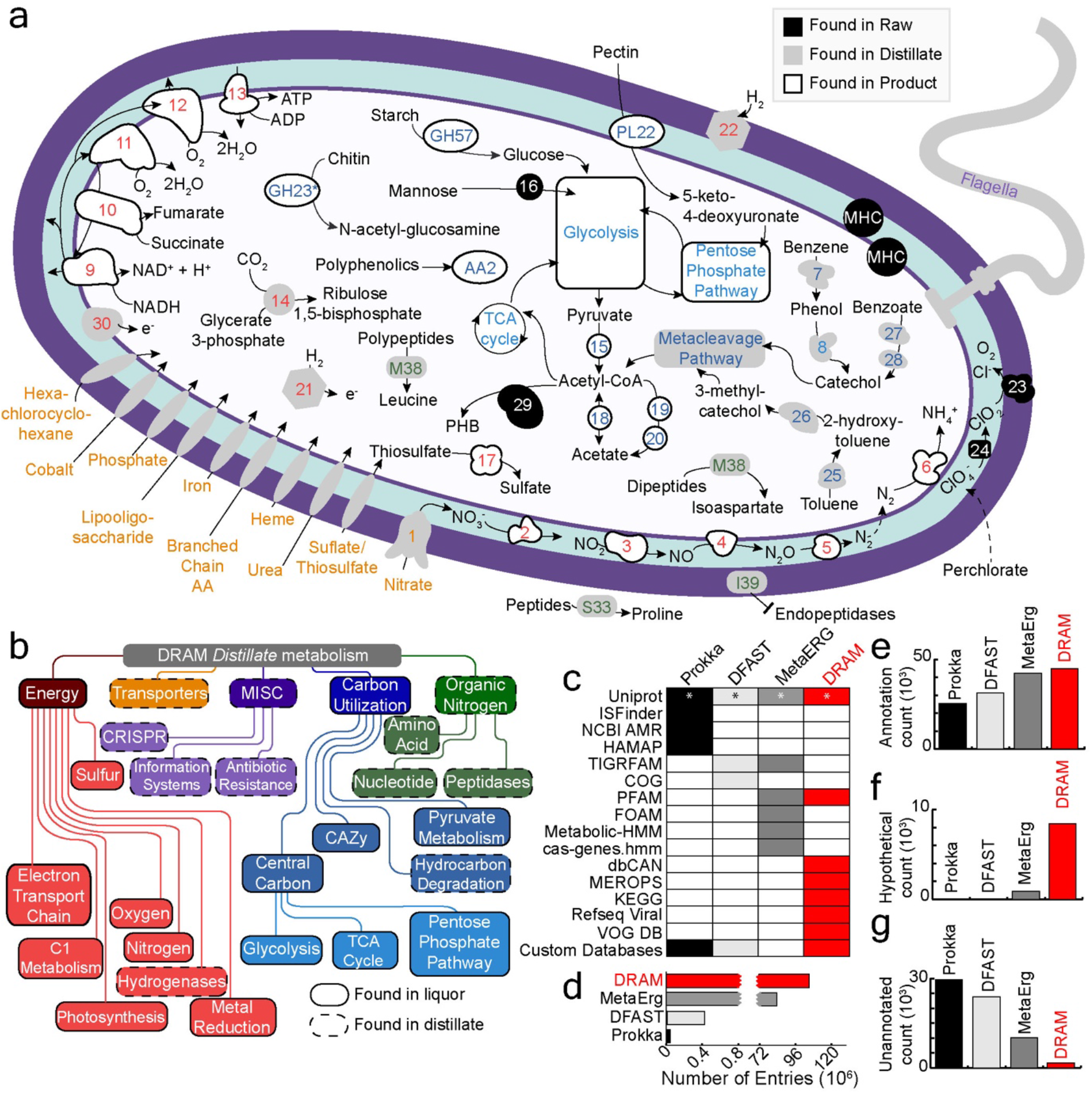
DRAM provides multiple levels of metabolic and structural information. **a** Genome cartoon of *Dechloromonas aromatica* RCB demonstrates the usability of DRAM to understand the potential metabolism of a genome. Putative enzymes are colored by location of information in DRAM’s outputs: *Raw* (black), *Distillate* (grey), and *Product* (white). Gene numbers, identifiers, or abbreviations are colored according to metabolic categories outlined in (**b**) and detailed in **Supplementary File 4**. Genes with an asterisk had an unidentified localization by PSORTb (102). **b** Flow chart shows the metabolisms from DRAM’s *Distillate. Distillate* provides five major categories of metabolism: energy, transporters, miscellaneous (MISC), carbon utilization, and organic nitrogen. Each major category contains subcategories, with outlines denoting location of information within *Distillate* and *Product*. **c** Heatmap shows presence (colored) and absence (white) of databases used in comparable annotators to DRAM. Annotators are colored consistently in a-e, with Prokka (30) in black, DFAST (31) in light grey, MetaErg (32) in dark grey, and DRAM in red. Barcharts in **d-g** show database size (**d**), as well as number of annotated (**e**), hypotheticals (**f**), and unannotated (**g**) genes assigned by each annotator when analyzing *in silico* soil community. See methods for definitions of annotated, hypothetical, and unannotated genes, relative to each annotator.

The *Distillate* output is further distilled to the *Product*, an HTML file displaying a heatmap (**Supplementary File 3**), created using Altair (52), as well as a corresponding data table. The *Product* has three primary parts: pathway coverage (e.g. glycolysis), electron transport chain component completion (e.g. NADH dehydrogenase), and presence of specific functions (e.g. mcrA, methanogenesis). The pathways selected for completion analysis were chosen because of their central role in metabolism. Pathway coverage is measured using the structure of KEGG (41) modules. Modules are broken up into steps and then each step is divided into paths. Paths can be additionally subdivided into substeps with subpaths. Coverage is given as the percent of steps with at least one gene present, substeps and subpaths are considered (**Supplementary Figure 2a**). This requires that at least one subunit of each gene in the pathway to be present. Electron transport chain component completion is measured similarly. Modules are represented as directed networks where KOs are nodes and outgoing edges connect to the next KO in the module. Completion is the percent coverage of the path through the network with the largest percentage of genes present (**Supplementary Figure 2b**). Function presence is measured based on the presence of genes with a set of identifiers. The gene sets were made via expert-guided, automatic curation of specific metabolisms (See Supplementary Text, section Interpreting results from DRAM and DRAM-v). Some functions require the presence of a single gene while others only require one or more annotations from sets of genes to be present (**Supplementary Figure 2c**). Specifics of the logic behind pathway completion, subunit completion, and specific functional potential calls are detailed in the Supplementary Text (section DRAM pathways and enzyme modularity completion).

### Benchmarking DRAM against commonly used annotators

In order to compare the performance in terms of runtime, memory usage and annotation coverage we compared DRAM to other commonly used genome or MAG annotation tools including Prokka (30), (v1.14.0), DFAST (31) (v1.2.3), and MetaErg (32) (v1.2.0) using three separate datasets: (i) *E. coli* strain K-12 MG1655, (ii) an *in silico* soil community we created (15 phylogenetically and metabolically distinct genomes from isolate and uncultivated Archaea and Bacteria), and (iii) a set of 76 MAGs generated from the largest HMP1(53) fecal metagenome (described below).

To compare annotation database size of each tool (Prokka, DFAST, and MetaErg) to DRAM, we counted the entries of each database used by default for each tool (**Figure 2cd, Supplementary File 4**). Specifically, for BLAST-based searches, the number of FASTA entries were counted for a given database, and for HMM-based searches, the number of model entries were counted for a given database.

To evaluate the annotation recovery by each tool, we compared the number of annotated, hypothetical, and unannotated genes assigned by each annotation tool to an *in silico* soil community and a set of MAGs generated from the largest HMP fecal metagenome (**Figure 2e-g**). A gene was considered *annotated* in DRAM if it had at least one annotation from KEGG (41), UniRef90 (42), MEROPs (43), Pfam (46) or dbCAN that was not “hypothetical”, “uncharacterized” or “domain of unknown function” gene. A gene is defined as *hypothetical* in DRAM if hits for a gene lacked defined annotation, and at least one of the annotations from KEGG (41), UniRef90 (42), MEROPs (43), Pfam (46) and dbCAN were “hypothetical”, “uncharacterized” or “domain of unknown function”. A gene was defined as *unannotated* in DRAM if no annotation was assigned from KEGG (41), UniRef90 (42), MEROPs (43), Pfam (46) or dbCAN (48). This is in contrast to other annotators, like Prokka (30) and DFAST (31) that remove many to all hypothetical genes from their databases and subsequently all genes are called as hypothetical, even genes that lack an annotation. Since these programs mask conserved hypothetical genes, the user loses the ability for broader biological context and further non-homology based discovery of protein function. In our performance analyses we considered DFAST and Prokka hypothetical labels as unannotated, as it was not possible to discern the difference between a gene that had no representatives in a database (unannotated) and a gene that had best hits to hypothetical genes in other organisms that were annotated in the database (hypothetical). In MetaErg (32), a gene was considered *unannotated* if in the master tab separated table there was no Swiss-Prot (54), TIGRFAM or Pfam (46) description. In MetaErg, a gene was considered *hypothetical* if hits lacked a defined annotation, and had at least one annotation from Swiss-Prot (54), TIGRFAM and Pfam (46) that contained “hypothetical”, “uncharacterized” or “domain of unknown function”.

Beyond differences in definition, we note that the summation of annotated, hypothetical, and unannotated genes is different for each tool due to the use of different gene callers or different filters on called genes, despite using the same input file (**Supplementary File 4**). Specifically, Prokka (30), MetaErg (32), and DRAM use Prodigal to call genes, while DFAST (31) uses MetaGeneAnnotator (55). But compared to DRAM, Prokka (30) filters out called genes that overlap with any RNA feature or CRISPR spacer cassette, while MetaErg (32) filters out all called genes <180 nucleotides. Default parameters were used for all annotation tools except for DRAM, which employed the --use_uniref flag to use UniRef to maximize the annotation recovery.

To measure speed and memory usage the three test sets were used with each annotation tool. All tools were run with default parameters. Each dataset and tool combination was run four times on the same machine using 10 Intel(R) Xeon(R) Gold 5118 CPU @ 2.30GHz processors. Average and standard deviations of run time and the maximum memory usage were reported. Performance data is reported in **Supplementary Figure 3a-c,** and **Supplementary File 4**.

The unit of annotation in DRAM is at the level of the gene, thus the number of genes (and not the number of genomes) in a dataset is the primary factor in determining runtime. In other words, assuming the same number of genes in the dataset, there would be no run time difference between the DRAM annotation of 100 unbinned, deeply sequenced, assembled metagenome samples and 10,000 binned, partial MAGs. For the datasets reported here, the gene numbers are 55,040 for a “mock” soil community and 143,551 for 76 MAGs assembled and binned from a HMP fecal metagenome, with the average run times for these data listed in **Supplementary Figure 3b**. To demonstrate scalability of DRAM, we also included the DRAM annotations for one of the largest MAG studies from a single ecosystem (21), with annotations provided for 2,535 MAGs (and including 6,273,162 total genes across the dataset) (https://zenodo.org/record/3777237). Summarizing, DRAM is scalable to an unlimited number of genes, however run time will be increased based on the number of genes annotated. In terms of the *Product* output, DRAM is not limited, but the *Product* heatmap is broken into sets of 1,000 genomes or metagenomes to facilitate effective visualization.

To address the accuracy of DRAM in recovering annotations for organisms with different levels of database representation, we used the most experimentally validated microbial genome, *E. coli* K12 MG1655 to annotate protein sequences with DRAM using different databases. We evaluated the 1) the full set of DRAM databases, 2) the full set of DRAM databases with all *Escherichia* genera removed, and 3) the full set of DRAM databases with all *Enterobacteriaceae* family members removed. The latter two databases (2 and 3) are meant to address assigning annotations of a microbial genome that may not have close representatives in the database (**Supplementary Figure 3d**).

### Selection of 15 Representative Soil Genomes for Annotation Benchmarking

To validate DRAM, we chose a set of phylogenetically diverse genomes from organisms with varying and known energy generating metabolisms. All genomes included in this analysis are from isolates, except for a member of the *Patescibacteria*, which was included to highlight the applicability of DRAM to Candidate Phyla Radiation (CPR) (**Supplementary File 4**). This dataset is not meant to represent an entire soil community, but rather was selected to highlight the metabolic repertoire (e.g. carbon, nitrogen, sulfur metabolisms) and phylogenetic divergence (different phyla across Bacteria and Archaea domains) commonly annotated in soil datasets.

Assembled nucleotide FASTA files for each genome or MAG were downloaded from NCBI or JGI-IMG. Genomes were annotated using DRAM.py annotate and summarized using DRAM.py distill (**Figure 3a-c**, **Supplementary Figure 1, Supplementary Files 3, 5**). Genomes were quality checked with checkM (51) and taxonomically classified using GTDB-Tk (v0.3.3) (24). Genome statistics and accession numbers are reported in **Supplementary File 4**.

**Figure 3:**
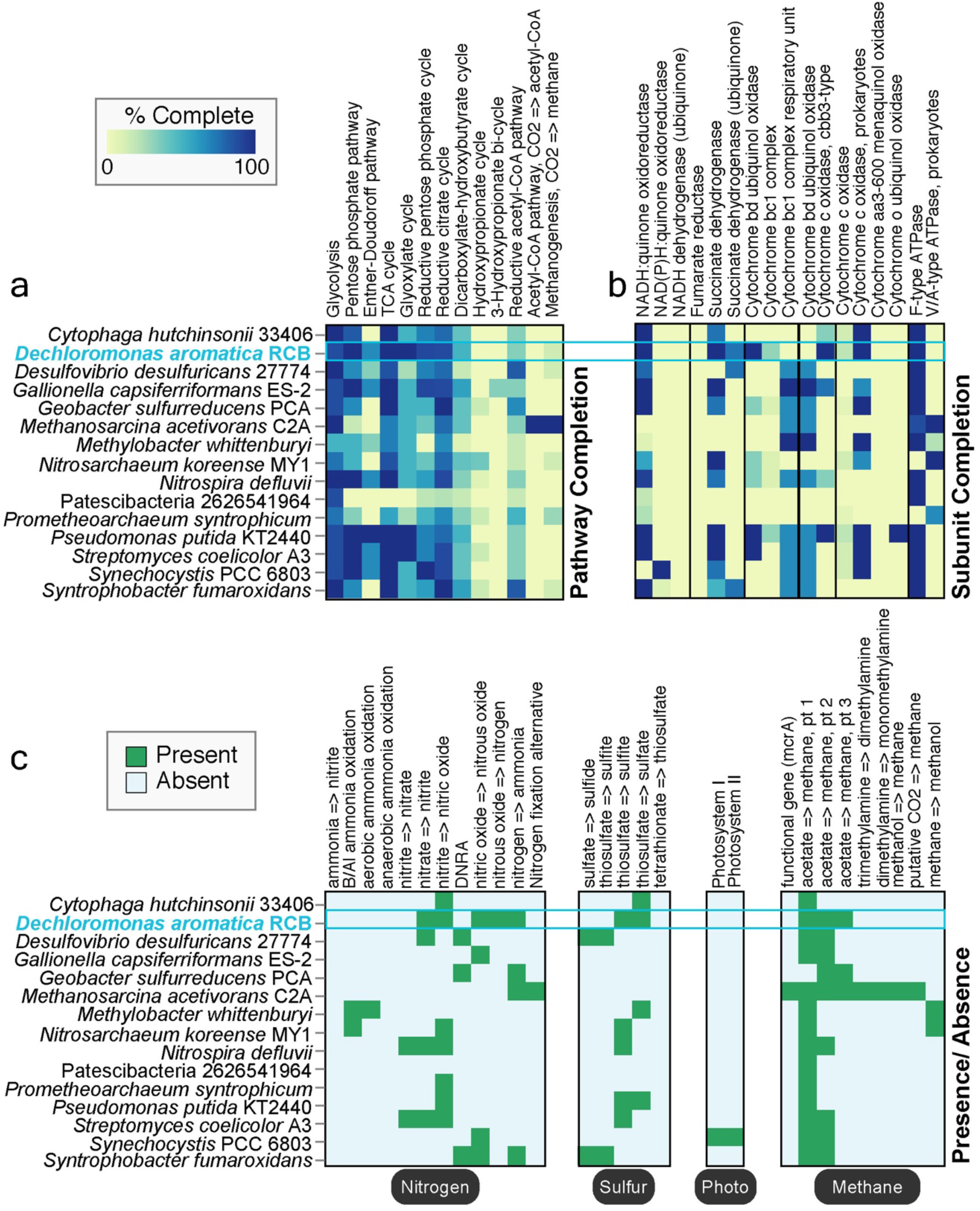
DRAM *Product* summarizes and visualizes ecosystem-relevant metabolisms across input genomes. Heatmaps in (**a**-**c**) were automatically generated by DRAM from the *Product* shown in **Supplementary File 3**. Sections of the heatmap are ordered to highlight information available in *Product*, including pathway completion (**a**), subunit completion (**b**), and presence/absence (**c**) data. Boxes colored by presence/absence in (**c**) represent 1-2 genes necessary to carry out a particular process. Hovering over the heatmap cells in the *Product’s* HTML outputs interactively reports the calculated percent completion among other information. *Dechloromonas aromatica* RCB is represented by a genome cartoon in **Figure 2a** and is highlighted in blue on the heatmaps.

### Human Gut Metagenome Samples Download and Processing

Forty-four human gut metagenomes were downloaded from the HMP data portal (https://portal.hmpdacc.org/) (**Supplementary File 4**) (53). All samples are from the HMP study (56) and are healthy adult subjects. All reads were trimmed for quality and filtered for host reads using bbtools suite (sourceforge.net/projects/bbmap/) (57). Samples were then assembled separately using IDBA-UD (58) using default parameters. The resulting assemblies were annotated using DRAM.py annotate and distilled using DRAM.py distill, resulting in 2,815,248 genes. To calculate coverage of genes, coverM (https://github.com/wwood/CoverM) was used in contig mode with the count measurement. These counts were then transformed to gene per million (GPM), which was calculated in the same manner as transcripts per million (TPM), with data reported in **Figure 4a-c**. To compare the variability of bulk level (*Distillate* categories) and substrate level categories across 44 human gut metagenomes, we calculated Bray-Curtis distances between all pairs of samples and used the Levene test to compare the variability of distances between annotations (**Supplementary Figure 4**).

**Figure 4:**
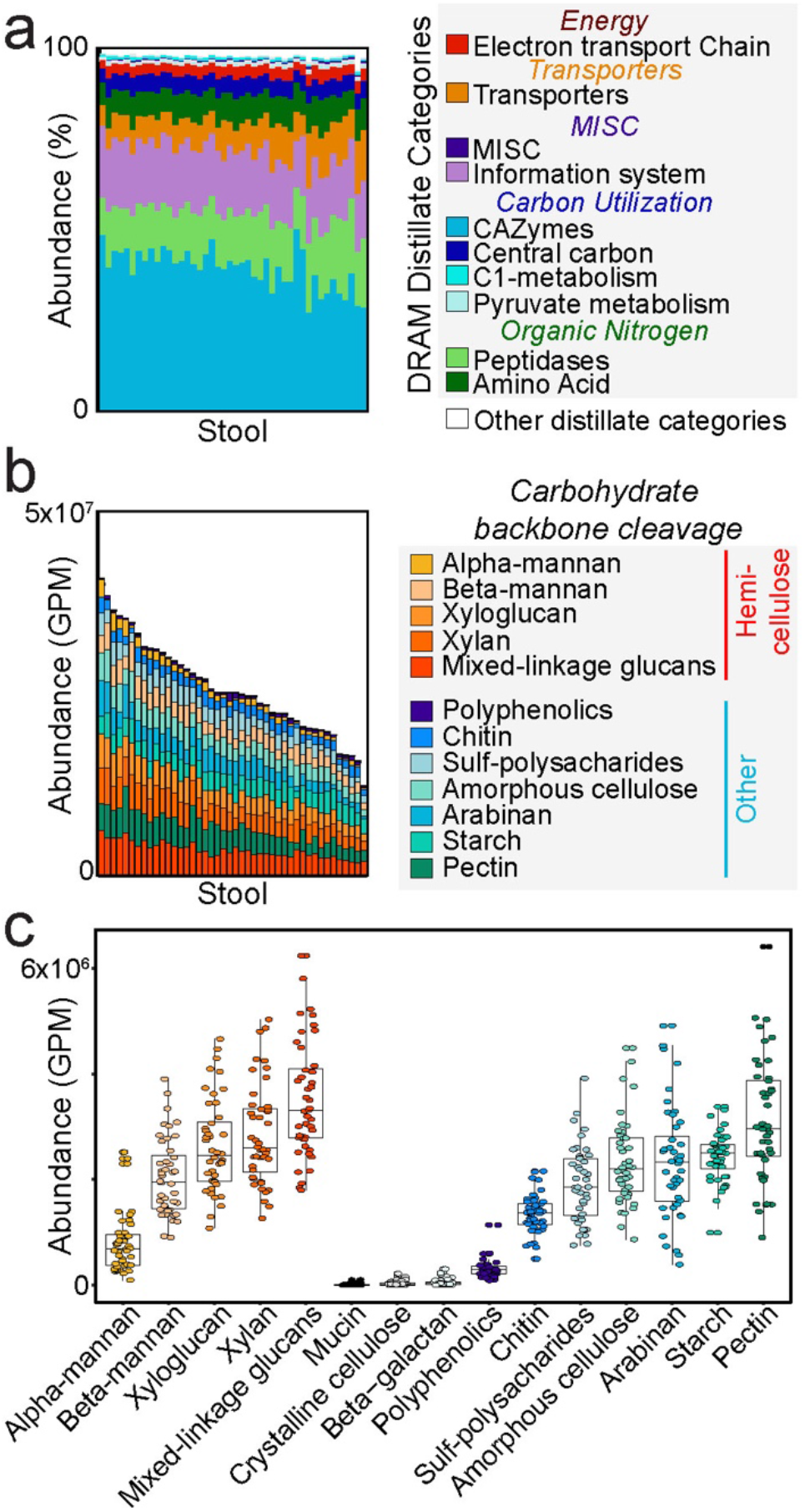
Substrate-resolved survey of carbon metabolism in the human gut. Bar charts represent normalized gene abundance or proportion of reads that mapped to each gene or gene category reported as relative abundance (%) or Gene Per Million (GPM). Reads came from previously (56) published healthy human stool metagenomes that were assembled and then annotated in DRAM (**a-c**). (**a**) Using a subset of 44 randomly selected metagenomes from (56), we profiled and annotated gene abundance patterns colored by DRAM’s *Distillate* categories and subcategories. (**b**) Using the same metagenomes and sample order as in (**b**), summary of CAZymes to broader substrate categories reveals differential abundance patterns across the cohort. (**c**) Data from (**b**) is graphed by carbohydrate substrates. Boxplots represent the median and one quartile deviation of CAZyme abundance, with each point representing a single person in the 44-member cohort. Putative substrates are ordered by class, then by mean abundance.

### Human Gut Metagenome for MAG Generation, Sample Download and Processing

To examine DRAMs ability to assign functionalities relevant to the human gut, we annotated MAGs present in a single Human Microbiome Project (53) sample. Raw reads from SRA accession number SRS019068 (the largest HMP metagenome collected to date, with 29 Gbp/sample) were downloaded from the NCBI Sequence Read Archive using wget (link: http://downloads.hmpdacc.org/dacc/hhs/genome/microbiome/wgs/analysis/hmwgsqc/v2/SRS019068.tar.bz2). Reads were trimmed for quality using sickle (https://github.com/najoshi/sickle) and subsequently assembled via IDBA-UD (58) using default parameters. Resulting scaffolds were binned using Metabat2 (59). We recovered 135 MAGs from this sample, that were dereplicated into 76 medium and high quality MAGs (60). Bins were quality checked with checkM (51), taxonomically classified using GTDB-Tk (v0.3.3) (24), and annotated and distilled using DRAM (**Figure 5**, **Supplementary Figure 5, Supplementary Files 1-2**). All assembly statistics and MAG statistics can be found in **Supplementary File 4**. To interrogate the importance of carbon metabolism in the human gut, the DRAM annotated CAZyme and SCFA production potential was profiled across the 76 medium and high quality MAGs using the DRAM *Distill* function. MAGs were clustered using hierarchal clustering via the hclust complete method in R (**Figure 5**).

**Figure 5:**
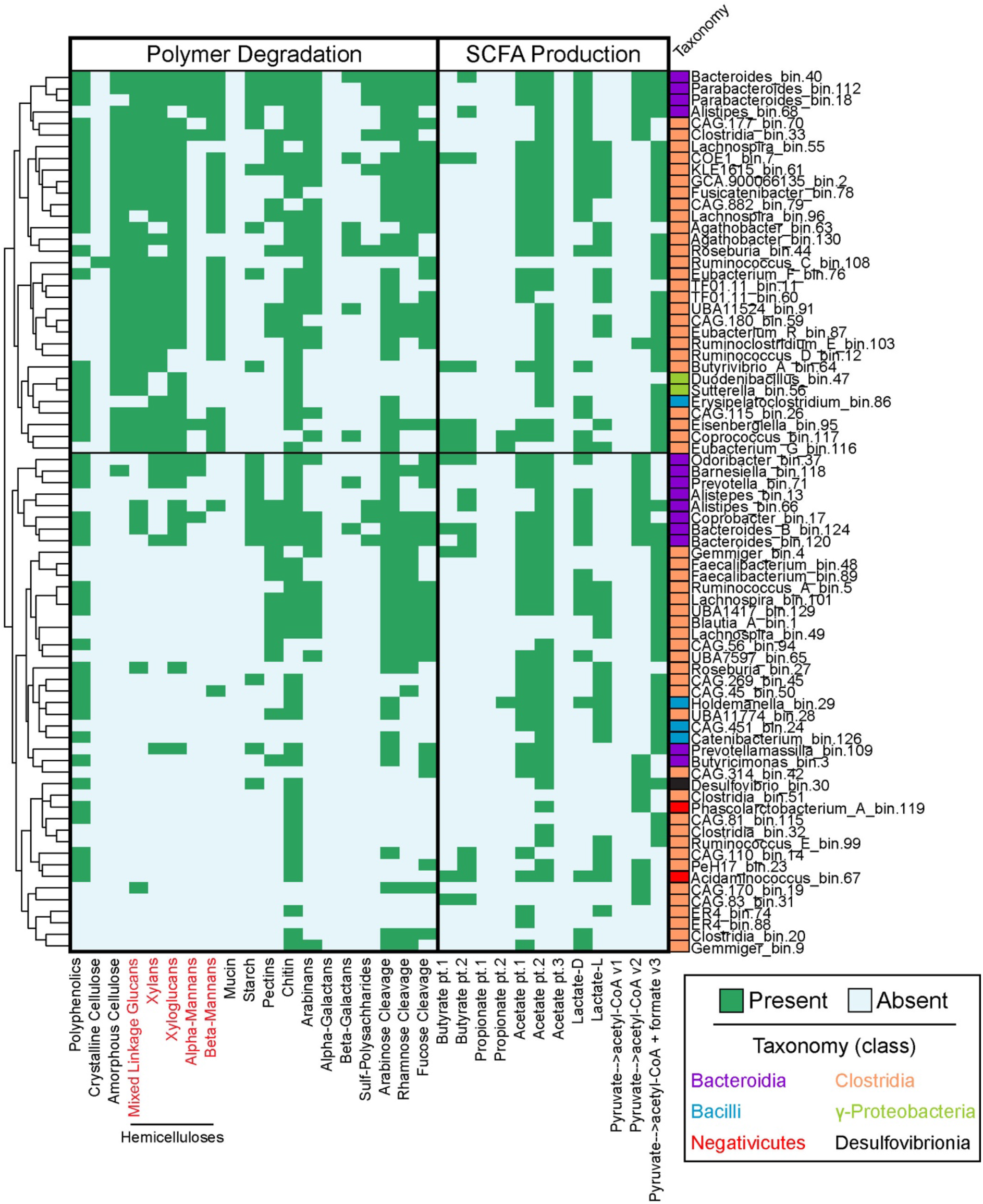
DRAM provides a metabolic inventory of microbial traits important in the human gut. Seventy-six medium and high-quality MAGs were reconstructed from a single HMP fecal metagenome. Taxonomy was assigned using GTDB-Tk (24), with colored boxes noting class and name noting genus. The presence (green) or absence (blue) of genes capable of catalyzing carbohydrate degradation or contributing to short chain fatty acid metabolism are reported in the heatmap. We note that the directionality of some of these SCFA conversions is difficult to infer from gene sequence alone. Genomes are clustered by gene presence and hemicellulose substrates are shown in red text.

### DRAM-v viral annotation and AMG prediction overview

The DRAM-v workflow to annotate vMAGs and predict potential AMGs is detailed in **Figures 1**, **6** and **Supplementary Figure 6**. DRAM-v uses VirSorter (61) outputs to find viral genomic (genomes or contigs) information in assembled metagenomic data. DRAM-v inputs must include a VirSorter (61) predicted vMAGs FASTA file and VIRSorter_affi -contigs.tab file. Each vMAG is processed independently using the same pipeline as in DRAM, with the addition of a BLAST-type annotation against all viral proteins in NCBI RefSeq. All database annotations in the DRAM-v results are merged into as single table as the *Raw* DRAM output.

**Figure 6:**
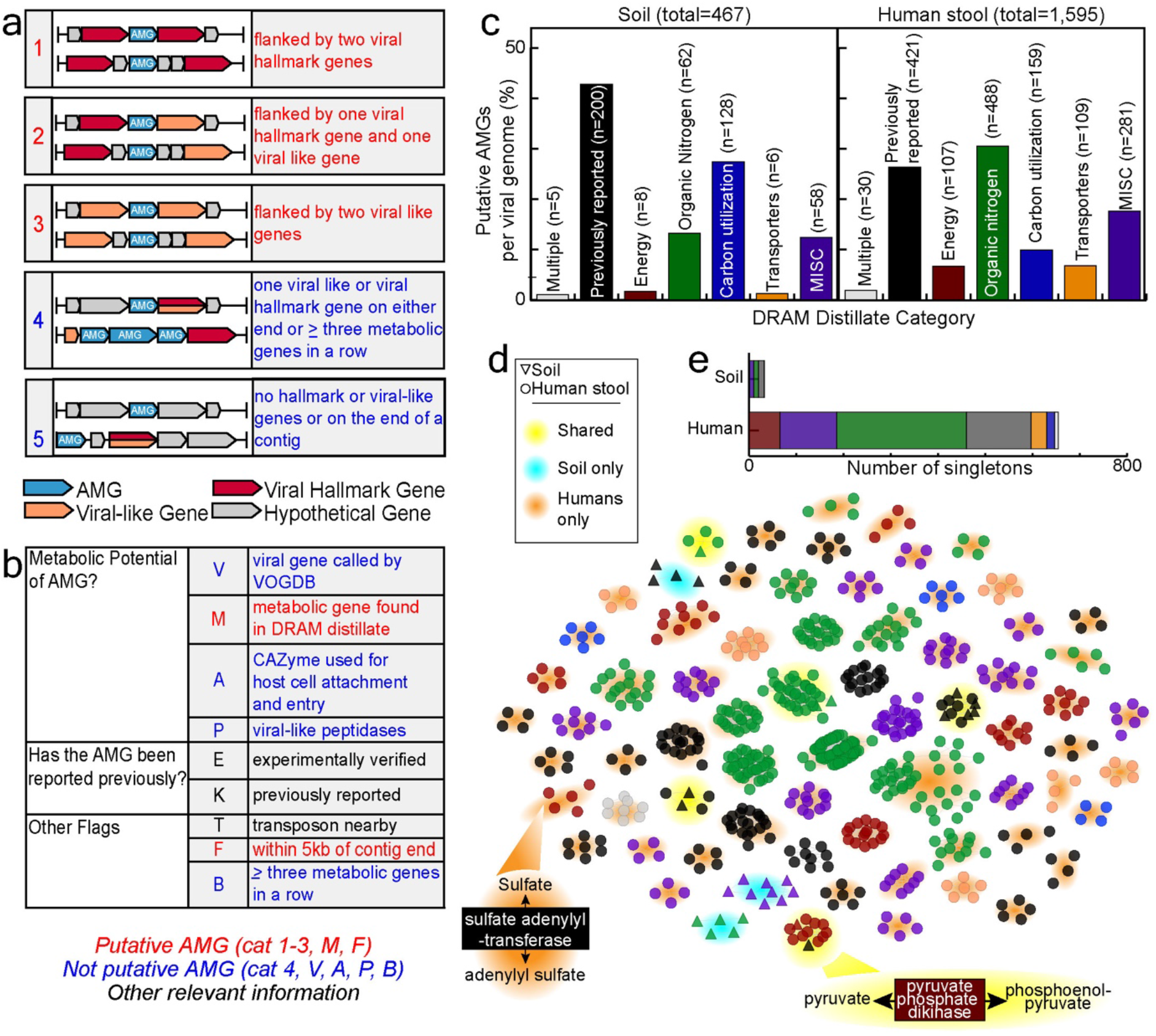
DRAM-v profiles putative AMGs in viral sequences. Description of DRAM-v’s rules for auxiliary (**a**) and flag (**b**) assignments. Auxiliary metabolic scores shown in (**a**) are determined by the location of a putative AMG on the contig relative to other viral hallmark or viral-like genes (determined by VirSorter (61)), with all scores being reported in the *Distillate*. Scores highlighted in red are considered high (1–2) or medium (3) confidence and thus the putative AMGs are also represented in the *Product*. Flags shown in (**b**) highlight important details about each putative AMG of which the user should be aware, all being reported in the *Raw*. Putative AMGs with a confidence score 1-3 and a metabolic flag (flag “M”; highlighted in red) are included in the *Distillate* and *Product*, unless flags in blue are reported. Flags in black do not decide the inclusion of a putative AMG. (**c**) Bar graph displaying putative AMGs recovered by DRAM-v from metagenomic files (soil metagenomes (14), left; 44 fecal metagenomes from the HMP (56), right) and categorized by the *Distillate* metabolic category: Carbon Utilization, Energy, Organic Nitrogen, Transporters and MISC. Putative AMGs labeled as “multiple” refer to genes that occur in multiple DRAM *Distillate* categories (e.g. transporters for organic nitrogen) and AMGs that are labeled as previously reported are in the viral AMG database compiled here. (**d**) Sequence similarity network (66) of all AMGs with an auxiliary score of 1-3 recovered from soil and human stool metagenomes. Nodes are connected by an edge (line) if the pairwise amino acid sequence identity is >80% (see **Methods**). Only clusters of >5 members are shown. Nodes are colored by the *Distillate* category defined in (**c**), while node shape denotes soil or human stool. Back highlighting denotes if the cluster contains both soil and human stool nodes (shared), soil nodes only, or human stool nodes only. Specific AMGs highlighted in the text are shown. (**e**) Stacked bar chart shows the number of singletons (AMGs that do not align by at least 80% to another recovered AMG) in each sample type, with bars colored by DRAM-v’s *Distillate* category.

After the annotation step, auxiliary scores are assigned to each gene. The auxiliary scores are on a scale from 1 to 5, and provide the user with confidence that a gene is on a vMAG (and not contaminating source). Here a score of 1 represents a gene that is confidently virally encoded and a score of 4 or 5 represents a gene that users should take caution in treating as a viral gene. These scores are based on previous manually curated data provided in **Supplementary File 4**. Auxiliary scores are assigned based on DRAM mining the category of flanking viral protein clusters from the VIRSorter _affi-contigs.tab file (**Figure 6a**). A gene is given an auxiliary score of 1 if there is at least one hallmark gene on both the left and right flanks, indicating the gene is likely viral. An auxiliary score of 2 is assigned when the gene has a viral hallmark gene on one flank and a viral-like gene on the other flank. An auxiliary score of 3 is assigned to genes that have a viral-like gene on both flanks. An auxiliary score of 4 is given to genes with either a viral-like or hallmark gene on one flank and no viral-like or hallmark gene on the other flank, indicating the possibility that the non-viral supported flank could be the beginning of microbial genome content and thus not an AMG. An auxiliary score of 4 is also given to genes that are part of a stretch with three or more adjacent genes with non-viral metabolic function. An auxiliary score of 5 is given to genes on contigs with no viral-like or hallmark genes and genes on the end of contigs.

Next, various flags that highlight the metabolic potential of a gene and/or qualify the confidence in a gene being viral are assigned (**Figure 6b**). The “viral” flag (V) is assigned when the gene has been associated with a VOGDB identifier with the replication or structure categories. The “metabolism” flag (M) is assigned if the gene has been assigned an identifier present in DRAM’s *Distillate*. The “known AMG” flag (K) is assigned when the gene has been annotated with a database identifier representing a function from a previously identified AMG in the literature. The “experimentally verified” flag (E) is similar to the (K) flag, but the AMG has to be an experimentally verified AMG in a previous study, meaning it has been shown in a host to provide a specific function (e.g. psbA photosystem II gene for photosynthesis (62, 63)). Both the (K) and (E) flags are called based on an expert-curated AMG database composed of 257 and 12 genes, respectively. The “attachment” flag (A) is given when the gene, while metabolic has been given identifiers associated with viral host attachment and entry (as is the case with many CAZymes). The viral “peptidase” flag (P) is similar to the (A) flag but when the gene is given identifiers that are peptidases previously identified as potentially-viral using, not AMGs, based on the distribution of peptidase families provided in the MEROPS (43) database. The “near the end of the contig” flag (F) is given when the gene is within 5,000 bases of the end of a contig, signifying that the user should confirm viral genes surrounding the putative AMG, as there is less gene content to surrounding the putative AMG. The “transposon” flag (T) is given when the gene is on a contig that contains a transposon, highlighting to the user that this contig requires further inspection as it may be a non-viral mobile genetic element (64, 65) (**Figure 6b**). The “B” flag is given to genes within a set of three or more consecutive genes assigned a metabolism flag “M”, signifying that this gene may not be an AMG and instead located in a stretch of non-viral genes (**Figure 6b**). Specifics of the logic behind the AMG flags (e.g. (P), (A), (B) flags) is detailed in the **Supplementary Text and Supplementary File 4**. In summary, DRAM-v flags automate expert curation of AMGs, with the intention to provide the user with known AMG reference sequences, indicate to the user viral genes that should not be considered AMGs, and cue the user to genes that require additional curation before reporting.

The distillation of DRAM-v annotations is based on the detection of potential AMGs. By default, a gene is considered a potential AMG if the auxiliary score is less than 4, the gene has been assigned an (M) flag, and has not been assigned as a peptidase or CAZyme involved in viral entry or metabolism (P or A flag), as a homolog to a VOGDB identifier associated with viral replication or structure (V flag), or the gene is not in a row of 3 metabolic genes (B flag) (**Figure 6**). The reported flags and minimum auxiliary score threshold can be changed by the user. All flags and scores were defined using experimentally validated AMGs (**Supplementary File 4**), and then were validated using a set of published AMGs from soil.

DRAM-v annotations are distilled to create a vMAG summary (DRAM-v *Distillate*) and a potential AMG summary (DRAM-v *Product*). The vMAG summary is a table with each contig and information about the contigs satisfying many MIUViG requirements^19^. Other information is also included in this output such as the VirSorter^17^ category of the virus, if the virus was circular, if the virus is a prophage, the number of genes in the virus, the number of strand switches along the contig, if a transposase is present on the contig, and the number of potential AMGs. We also summarize the potential AMGs giving the metabolic information associated with each AMG as found in *Distillate*. DRAM-v’s *Product* further summarizes the potential AMGs showing all vMAGs, the number of potential AMGs in each contig, and a heatmap of all possible *Distillate* categories to which each AMG (category 1-3, default) belongs.

### Retrieval and Processing of Emerson et al. Data

1,907 vMAGs reported by Emerson *et al*. (14) were retrieved from DDBJ/ENA/GenBank via the accession number QGNH00000000. These contigs were processed with VirSorter 1.0.3 (61) in virome decontamination mode to obtain categories and viral gene information necessary for DRAM-v. Viral sequences with viral categories 1 and 2 and prophage categories 4 and 5 retained (1,867 contigs). DRAM-v was then run with default parameters, and the *Distillate* table is reported in **Supplementary File 6** and the *Product* is in **Supplementary File 7**.

### Processing of HMP Viral Sequences

Viral sequences were identified in the assembled HMP metagenomes using VirSorter 1.0.3 (61) hosted on the CyVerse discovery environment. VirSorter (61) was run with default parameters using the ‘virome’ database and viral sequences with viral categories 1 and 2 and prophage categories 4 and 5 were retained (2,932 contigs). Resulting viral sequences were annotated using DRAM-v.py annotate (min_contig_size flag set to 10,000) and summarized using DRAM-v.py distill (**Supplementary File 8-9**). All viral genomes used or recovered in this study are reported in **Supplementary File 10**.

### Generation of AMG Sequence Similarity Network

To identify the AMGs shared across systems, sequence similarity networks were generated via the EFI-EST webtool (66) using putative AMGs recovered from soil (n=547) and stool (n=2,094) metagenomes via DRAM-v as the input. A minimum sequence length of 100 amino acids, no maximum length, and 80% amino acid identity was specified from initial edge values. Representative networks were generated and visualized in Cytoscape 3.7.2 (67). Edge scores were further refined and *Distillate* categories and system information were overlaid in Cytoscape (67). **Figure 6** contains the resulting network filtered to clusters >5.

### Virus host matching in a single HMP sample

For the single binned HMP sample (SRS019068), viral sequences were matched to host MAGs using the CRISPR Recognition Tool (68) plugin (version 1.2) in Geneious. To identify matches between viral protospacers and host CRISPR–Cas array spacers, we used BLASTn with an e-value cutoff of 1 × 10-5. All matches were manually confirmed by aligning sequences in Geneious, with zero mismatches allowed. There was one virus (scaffold_938) that had a CRISPR host match and a putative AMG (genes HMP1_viralSeqs_398_VIRSorter_scaffold_938-cat_2_58-HMP1_viralSeqs_398_VIRSorter_scaffold_938-cat_2_59), with details provided in the Supplementary Text, section *Integration of DRAM and DRAM-v to begin to infer virocell metabolism*.

### Adding metabolisms to DRAM

DRAM is a community resource, as such we welcome metabolism experts to help us build and refine metabolisms analyzed in DRAM. Visit this (<u>link</u>) to fill out the google form, your metabolism will be vetted, and you receive an email from our team.

## RESULTS

### Enhanced annotation and distillation of genome attributes with DRAM

Like the process of distillation, DRAM generates and summarizes gene annotations across genomes into three levels of refinement: (1) *Raw*, (2) *Distillate*, and (3) *Product* (**Figure 1**). The *Raw* is a synthesized annotation of all genes in a dataset across multiple databases, the *Distillate* assigns many of these genes to specific functional categories, and the *Product* visualizes the presence of key functional genes across genomes. Through this high-throughput distillation process, DRAM (**Figure 1a**), and the companion program DRAM-v (**Figure 1b**), annotates and organizes high volumes of microbial and viral genomic data, enabling users to discern metabolically relevant information from large amounts of assembled microbial and viral community sequencing information.

The *Raw* annotations provided by DRAM are a comprehensive inventory of multiple annotations from many databases. These *Raw* annotations are where most other annotators stop, with analyses in the DRAM *Distillate* and *Product* uniquely designed to expedite the functional and structural trait profiling within and across genomes (**Figure 2a**). In the *Distillate*, the DRAM *Raw* data is parsed into five categories and subsequent subcategories (**Figure 2b**). With the goal to standardize the reporting of genome quality across publications, the minimum suggested standards for reporting MAGs (25) are also summarized in the *Distillate*. Specifically, DRAM compiles the quantification of tRNAs, rRNAs, and genome size metrics (e.g. length, number of contigs) with user provided estimates of genome completeness, contamination (51), and genome taxonomy (24). This summation is synthesized into a quality metric for each genome that includes a rank of high, medium, or low quality based on established standards (25).

The *Product* is the most refined level of DRAM, and uses functional marker genes to infer broad metabolic descriptors of a genome. This summary of genes enables classification of the respiratory or fermentative metabolisms encoded in a genome, while also accounting for selected carbon metabolic pathways (**Figure 3, Supplementary File 3**). Moreover, completion estimates are calculated for electron transport chain complexes or pathways (**Figure 3**). We note these completion metrics are based on the percentage of genes recovered for unique subunits or physiological steps (**Figure 3a**), which is in contrast to analyses from other tools that recognize all non-redundant routes as equivalent (**Supplementary Figure 2**). This provides more accurate pathway completion estimates, as certain pathways are often underestimated when less physiologically refined approaches are used. The *Product* provides an interactive HTML heatmap that visualizes the presence of specific genes, including the gene identifiers which allow the user to link data across all DRAM levels (in the *Raw* and *Distillate*).

We recognize that DRAM is a first step in the annotation process, and thus the DRAM outputs are designed to make it convenient to export content at the gene, pathway, or genome level (e.g. FASTA or GenBank files). To help the user navigate the DRAM levels, we constructed a genome metabolic cartoon based on DRAM annotations of an isolate genome (*Dechloromonas aromatica* strain RCB) (**Figure 2a, Supplementary File 4**). We use this figure to illustrate where different genetic attributes reside in DRAM. Notably, DRAM has the ability to distill microbial metabolism for thousands of individual genomes simultaneously, which allows users to easily compare and identify patterns of functional partitioning within an entire microbial community.

### DRAM recovers more annotations compared to other assembly-based annotation software

We first compared the overall features of DRAM to common genome annotators or viewers (**Supplementary Table 1**), finding that published annotation systems often lack the ability to scale across thousands of genomes, visually summarize metabolism, or annotate virally encoded metabolic functions. Next to benchmark DRAM performance, we compared the DRAM database content and performance criteria to results from published MAG annotation tools (Prokka (30), DFAST (31), and MetaErg (32)), which are three commonly used pipelines for genome annotation with multi-genome files (**Supplementary Table 1**). To maximize annotation recovery, DRAM incorporates 7 different databases that provide functionally disparate, physiologically informative data (e.g. MEROPS (43), dbCAN2 (48)), rather than overlapping content (e.g. HAMAP, UniProt) (**Figure 2c**). Beyond just using more databases for annotation, DRAM also provides expert curation of this content (e.g. dbCAN2, MEROPS) (see Supplementary Information, Interpreting results from DRAM and DRAM-v). Moreover, for the UniProt database (69) shared across these annotators, DRAM uses the most comprehensive version (Uniref90 (42)) compared to other annotators that use a proprietarily culled version of the database resulting in 132-to 3,412-fold less entries. Summing all the databases used for each annotator, DRAM has millions more entries (from 21M to 104M) (**Figure 2d**, **Supplementary File 4**).

We next evaluated the annotation recovery of DRAM relative to published annotation tools by quantifying the number of annotated, hypothetical, and unannotated genes assigned by each tool (30–32) from an *in silico* soil community we created (15 phylogenetically and metabolically distinct genomes from isolate and uncultivated Archaea and Bacteria) (**Supplementary File 4**). Compared to the other annotators, for the *in silico* soil community, DRAM recovered 44,911 annotated genes, which was on par with MetaErg (32) (42,478 genes), but 1.4-1.8 times more than Prokka (30) and DFAST (31) (25,466 and 31,258 genes, respectively). Compared to other tools, DRAM better differentiates homologs with a hypothetical annotation from unannotated genes (see Methods, **Figure 2e-g, Supplementary Figure 3**). This increased identification of hypothetical annotations allows users to find homologs conserved in other organisms, providing hypotheses for gene function that can be further validated by experimental characterization (70). The reduction of unannotated genes is most notable for the Patescibacteria genome, a MAG from an uncultivated lineage in our *in silico* soil community. For this genome, DRAM produced 825 annotated, 362 hypothetical, and 7 unannotated genes, compared to 802 annotated, 11 hypothetical, and 342 unannotated genes output from the next closest annotator (32). Beyond increased annotation and hypothetical yield, DRAM also produced more meaningful annotations that can be readily incorporated into models, with DRAM recovering more EC numbers for this Patescibacteria genome compared to other tools (**Supplementary File 4**). To further test the performance of DRAM, we annotated the *E. coli* K-12 MG1655 genome using filtered versions of the KEGG Genes database to quantify precision and recall. Performance metrics were highest when the genes from the *E. coli* K-12 MG1655 genome were present in the database, but even when the entire genus of *Escherichia* was removed, performance remained high, with precision falling by 0.1% and recall falling by 0.8%, suggesting DRAM with default settings is relatively conservative and sacrifices recall for high levels of precision (**Supplementary Figure 3**).

We note, however, that this increased annotation quality and synthesis comes at expense of run time and potentially overall memory usage (depending on database selection), with genomes from the *in silico* soil community having an average complete annotation time (Raw, Distillate, Product) of 15 minutes per genome (**Supplementary Figure 3**). Unlike run time, memory usage is only minorly impacted by the number of genes analyzed (~1 MB per genome, (**Supplementary Figure 3**)), but is impacted by the database selection (especially UniRef90 (42)). For example, DRAM memory use doubled from running the same samples with (~200 GB) and without (~100 GB) UniRef90 (42). Thus, if memory usage or access to databases is limited, we provide the option to modify the DRAM databases (see Methods). In summary, DRAM is scalable to thousands of genomes albeit run time is impacted by number of genes analyzed. To demonstrate the scalability of DRAM, we annotated one of the largest MAG datasets from a single ecosystem (21), highlighting the ability of DRAM to summarize the metabolic potential of thousands of genomes at once (https://zenodo.org/record/3777237). Beyond annotation recovery and resolution, DRAM has more downstream functionalities and synthesis than other tools (**Supplementary Table 1**).

### DRAM profiles diverse metabolisms in an in silico soil community

To evaluate the capacity of DRAM to rapidly profile different metabolic regimes across genomes, we created an *in silico* soil community made up of phylogenetically distinct and metabolically versatile organisms (**Supplementary File 4**). For 13 of the 14 genomes with a cultivated representative in our *in silico* soil community, the findings from DRAM were consistent with prior broad-scale physiological classifications for each isolate (**Figure 3**). For a single genome in our dataset, a known ammonia oxidizing isolate that has not been reported to perform methane oxidation (*Nitrosoarcheaum koreense* MY1), DRAM reports the presence of a functional gene for methanotrophy (*pmoA*). We include this example to highlight how the well-documented sequence similarity between *amoA* for ammonia oxidation and *pmoA* for methane oxidation causes difficulty in reconciling proper function through homology based queries used in all multi-genome annotators today including Prokka, DFAST, and MetaErg (30–32, 71, 72). Consequently, DRAM is only a first step in identifying key functional genes, as subsequent non-homology based methods (e.g. phylogenetic analyses, protein modeling (73), gene synteny, Bayesian inference framework (74, 75)) or physiological or biochemical characterization are often required to validate findings from any homology-based annotator.

Within organisms reported to have the potential to respire (11/15 genomes), all were correctly identified in the DRAM *Product* by the presence of a complete NADH or NADPH dehydrogenase complex and a complete TCA pathway in the genome (**Figure 3ab**). The DRAM *Product* profiles the capacity to respire oxygen (e.g. *Pseudomonas putida*), nitrate (*Dechloromonas aromatica*), sulfate (*Desulfovibrio desulfuricans*), and others (**Figure 3c**). Additionally, photorespiration and methanogenesis are summarized in the *Distillate* and *Product*, exemplified by the photosynthetic *Synechocystis* sp. PCC 6803 and methanogenic *Methanosarcina acetivorans* (**Figure 3c**). Using two model genomes that encode the capacity for obligate fermentation (3, 76), one cultivated (*Candidatus Prometheoarchaeum syntrophicum* strain MK-D1) and one MAG from an uncultivated *Patescibacteria* (24) (also Parcubacteria genome GW2011_GWF2 (3)), we show that DRAM reasonably profiles carbon use and fermentation products. The value of using enzyme complex completion to reduce misannotations is demonstrated (**Supplementary Figure 2**), as the partial completion (3 genes) of the multi-subunit NADH dehydrogenase is not due to a complete complex I, but rather the presence of a trimeric hydrogenase common in obligate fermenters (3, 77). These hydrogenases are further annotated in detail by their type and function in the *Distillate*. In summary, the *Product* accurately assigns broad biogeochemical roles to this mock soil community, demonstrating the breadth of metabolisms that can be visualized and rapidly analyzed across multiple genomes from isolate and metagenome sources.

### DRAM uncovers personalized, substrate specific carbohydrate utilization profiles in the human gut

While mock communities like our prior soil community are commonly used for software performance criteria, they typically represent simpler communities than what is found in real-world samples. To demonstrate the feasibility of DRAM to apply to contemporary, complex, authentic samples, we analyzed the metabolic features of 44 HMP unbinned fecal metagenome samples. These samples had an average of 6.1 Gbp (with a maximum of 17 Gbp) per sample, consistent with or exceeding the average sequencing depth per sample reported in recent human gut studies in the last two years (56, 78, 79) (**Supplementary File 4**). These HMP metagenomes were selected from a landmark study that used COG defined categories to describe the microbially encoded traits in a cohort of healthy humans (56). Using broad process level categories (e.g. central carbohydrate metabolism), it was concluded in this publication (56) that microbial functional gene profiles were consistent across humans. DRAM is also able to evaluate gene content at broad categories, showing that CAZymes and peptidases are most prevalent in these datasets (**Figure 4a**). From this data, we hypothesized that increasing the resolution to the substrate level would reveal more personalized phenotypic patterns that were previously undefined in this cohort. To test this hypothesis for carbohydrate use, we used DRAM to classify bacterial and archaeal glycoside hydrolases, polysaccharide lyases, and enzymes with auxiliary activities related to carbohydrate-active enzymes (CAZymes (48)). DRAM then parsed this information, producing a microbial substrate utilization profile for the gut microbial community in each human. We note, that this assignment is not unambiguous as some CAZymes are promiscuous for multiple substrates (79), a functionality DRAM accounts for in the *Distillate* and *Product* (**Supplementary Figure 2, Supplementary File 4**). Consistent with our hypothesis, carbohydrate substrate use profiles predicted by DRAM were more variable than bulk level DRAM *Distillate* annotations across humans (**Supplementary Figure 4**). This more resolved annotation showed a 3-fold difference in CAZyme gene relative abundance across the cohort (**Figure 4bc**). In summary, using more resolved annotations will likely reveal that the gut gene content is not as stable as historically perceived (56). Specifically, CAZymes with the capacity to degrade hemicellulose components had the greatest mean abundance (3×10^7^ GPM), pectin was the most variable (7-fold change), and mucin had the most variable detection (only in 50% of cohort) (**Figure 4d**). Interestingly, the dominance of hemicellulose and the variability of pectin is reflective of the western diet, which is high in the consumption of cereal grains and not uniform in the consumption of fruit and vegetables (80–82). Our findings illustrate how DRAM substrate inventories could uncover linkages between gut microbiota gene content and host lifestyle or host genetics. Similarly, shifts in carbohydrate use patterns have been shown to be predictive of human health and disease (83, 84), thus this added level of annotation refinement provided by DRAM in an easy-to-understand format makes it possible to resolve biochemical transformations occluded by bulk level annotations.

### MAG profiles for utilization of specific organic carbon and nitrogen substrates generated by DRAM

To show that DRAM can not only profile the function of an entire microbial community, but can also parse metabolisms to specific genomes within this community, we assembled the largest (29 Gbp) publicly available Human Microbiome Project (HMP) fecal metagenome. We recovered 135 MAGs, of which 75 were medium quality and 1 was high-quality as assessed by DRAM. The taxonomic assignment of these MAGs according to DRAM taxonomy summary from GTDB (24) was predominantly Firmicutes and Bacteroidota, with rare members affiliated with the Proteobacteria and Desulfobacterota (**Supplementary Figure 7**). The taxonomic identity of the MAGs we recovered using this binning approach (previously the sample was unbinned), are similar to the membership reported in the healthy, western human gut (85), indicating this sample can serve as a reasonable representative to demonstrate DRAMs annotation capabilities of gut MAGs.

In the mammalian gut, beyond the digestion of carbohydrates with CAZymes, microorganisms also play critical roles in processing dietary protein into amino acids via peptidases (86) and producing short chain fatty acids for host energy as a fermentation byproduct (87). From these 76 HMP genomes, DRAM identified 7,197 and 5,471 CAZymes and peptidases, respectively (**Figure 5, Supplementary Figure 5, 8, Supplementary Files 1-2**). The capacity to degrade chitin was the most widely encoded (81%) across the genomes, a capacity reported to increase during gut inflammation (88). We also show that the capacity to cleave glutamate from proteinaceous compounds is the most commonly detected in our genomes, likely reflecting high concentrations of this amino acid in the gut (89). The substrate resolution provided by DRAM will enable more detailed analysis of microbial community inputs and outputs relevant to understanding the gut microbiomes impact on human health and disease (9, 84, 90, 91).

Given the importance of SCFA metabolism in the gut ecosystem, we show DRAMs capability to profile these metabolisms. It is no surprise that this capability is widely encoded by phylogenetically distinct genomes. Among the 76 HMP MAGs, the potential for acetate production was the most widely encoded, while propionate production potential was the least prevalent. The gene relative abundance reflects reported metabolite concentrations in the mouse and human gut (87, 92). Collectively, these results show how outputs of DRAM can be used to establish hypotheses for carbohydrate utilization trophic networks, where metabolic interactions can be considered simultaneously, rather than oversimplified into pairwise interactions (93). Moreover, by making it easier to assay substrate and energy regimes, it is our hope that DRAM can assist in development of designer cultivation strategies and the generation of synthetic communities for desired degradation outcomes.

### DRAM-v, a companion tool to systematically automate identification of viral auxiliary metabolic genes

Viruses are most often thought of as agents of lysis – impacting microbial community dynamics and resource landscapes. However, viruses can also impact microbial functioning and biogeochemical cycling via encoding and expressing Auxiliary Metabolic Genes (AMGs) (94) that directly alter host metabolisms during infection. To date, AMG annotation from viral isolates (62, 95) and metagenomic files (14, 15) has not scaled with the rate of viral genome discovery. Further, there are now numerous examples of metabolic genes in “viromes” that are more likely to be microbial DNA contamination (39), which is even a greater concern in metagenomic files where the resultant viruses can include prophages whose ends are challenging to delineate (61, 96). To automate the identification of putative AMGs, we sought to complement DRAM with a companion tool, DRAM-v, that (i) leverages DRAM’s functional annotation capabilities to describe metabolic genes, and (ii) applies a systematic scoring metric to assess the confidence for whether those metabolic genes were within *bona fide* viral contigs and not microbial (**Figure 1b, Supplementary Figure 6**). To demonstrate how these scoring metrics and ranks come together in our AMG annotation, see the example output files (Supplementary Files 6-9).

For each gene on a viral contig that DRAM-v has annotated as metabolic, we developed an auxiliary score, from 1 to 5 (1 being most confident), to denote the likelihood that the gene belongs to a viral genome rather than a degraded prophage region or a poorly defined viral genome boundary (**Figure 6a**). Because viral resources remain underdeveloped and several ambiguities can remain for some ‘hits’ even after these auxiliary scores are applied, DRAM-v uses flags to help the user quickly see where possible AMGs have been experimentally verified or previously reported. DRAM-v also flags users to the probability of a gene being involved in viral benefit rather than enhancing host metabolic function (e.g. certain peptidases and CAZymes are used for viral host cell entry (**Figure 6b**). DRAM-v, like DRAM, also groups viral genes into functional categories, provides quality reporting standards for viral contigs (27), and visualizes the predicted high- and medium-ranked AMGs (auxiliary scores 1-3) in the *Product*. DRAM-v and the AMG scoring system established here make it possible to rapidly identify viruses capable of augmenting host metabolism.

To benchmark the precision of DRAM-v, we reannotated viral contigs from a soil metagenomic file that our team had manually curated for glycoside hydrolase AMGs in a previous study (14). In that study, we reported 14 possible glycoside hydrolase AMGs from over 66,000 predicted viral proteins on viral contigs >10 kbp (14). Reannotating this file using DRAM-v, we recovered 100% of these AMGs according to DRAM’s defined metrics. Moreover, we recovered an additional 453 genes that were ranked with high (auxiliary scores 1, 2) or medium (auxiliary score 3) AMG confidence (**Supplementary File 6-7**). Because DRAM expands the metabolic repertoire and the speed at which metabolisms could be inventoried across hundreds of viral contigs, we were able to increase the AMG recovery by 32-fold. Our DRAM-v findings show that soil viral genomes encode AMGs that could play roles in host energy generation (2%), carbohydrate utilization (27%), and organic nitrogen transformation (13%) (**Figure 6c**). Moreover, 42% of the putative AMGs had been previously reported in other files.

### DRAM-v uncovers conserved and unique AMGs across ecosystems

We harnessed the automation and functional categorization power of DRAM-v to understand how viral AMG diversity varies across ecosystems. To that end, we recovered 2,932 viral contigs, containing 1,595 putative AMGs from the 44 HMP metagenome samples discussed above (**Figure 4, Supplementary Files 8-10**) and compared these AMGs to the 467 putative AMGs that we recovered from the soil metagenomes discussed above (**Figure 6c**). The majority of the HMP AMGs had putative roles in energy generation (7%), carbon utilization (10%), and organic nitrogen transformations (30%). The human gut is nitrogen limited (97), which may explain why putative AMGs for organic nitrogen transformations were the most well represented (**Figure 6c**). Specifically, the majority of the organic nitrogen AMGs we identified in the gut were likely involved in augmenting microbial host amino acid synthesis and degradation capacities. AMGs for tyrosine (EC 1.3.1.12, prephenate dehydrogenase) and lysine (EC 4.1.1.20, diaminopimelate decarboxylase) synthesis were of particular interest as they were uniquely encoded in specific phage genomes and had high quality auxiliary scores (**Supplementary File 8**). These AMGs could be valuable for their microbial hosts, given that increased gene copy number in these pathways was shown to enhance microbial growth (98). Moreover, synthesis of these branched and aromatic amino acids is costly for the microbial host and these compounds are absorbed by gut epithelial cells (99), thus there are clear advantages for hosts that can rapidly synthesize these scarce resources.

To directly compare soil and gut viral AMGs, AMG counts were normalized to the number of viral contigs in each file. Overall, stool viruses encoded more putative AMGs compared to soil viruses. These soil AMGs were mostly associated with carbon utilization, while gut AMGs were more linked to organic nitrogen transformations (**Figure 6c**). To identify shared and unique AMGs across these two files, we built an amino acid sequence similarity network of all the recovered AMGs (**Figure 6d**). Notably, the majority of putative soil AMGs, particularly CAZymes, do not share sequence similarity with gut-derived AMGs (**Figure 6e**). AMGs shared between soil and human stool are related to organic nitrogen or energy metabolisms.

AMGs within energy categories were of particular interest, as these genes may increase the copy number resulting in greater activity, or expand the metabolic repertoire of the host (38). For example, sulfate adenylyl transferase identified in soils is a key gene for sulfur assimilation and dissimilation, while pyruvate phosphate dikinase, a gene to promote the metabolism of this key central carbon metabolite, was shared by both soil and human gut ecosystems (**Figure 6d**). The conservation and uniqueness of these AMGs across ecosystems hints at more universal and environmentally tuned roles that virus may play in modulating their host and surrounding environment (**Supplementary Figure 9-10, Supplementary File 4**). We note that while DRAM is an important first step in the rapid and uniform detection of viral AMGs, contextualizing the physiological and biochemical role of AMGs requires additional analyses (14).

## DISCUSSION

DRAM provides a scalable and automated method for annotating features of assembled microbial and viral genomic content from cultivated or environmental sequencing efforts. This unparalleled annotation tool makes inferring metabolism from genomic content accessible. Here we show that DRAM is a critical, first step in annotating functional traits encoded by the microbiome (100). To facilitate further recommended curation, DRAM provides outputs in formats interoperable with downstream phylogenetic approaches (101), membrane localization analyses (102), visualization by genome browsers (103), and protein-structural modelling (73). DRAM annotations, like all homology-based genome annotation tools commonly used today, are reliant on the content in underlying databases. We show here that the variety of databases used in DRAM contributes to enhanced annotation recovery. Moreover, looking to the future, we built the DRAM platform to be robust, and with the capability to ingest non-homology based annotations as well.

Beyond the content in databases, it is our hope that DRAM can ease the dissemination of emerging metabolisms and biochemistry, offering a community resource to rapidly assimilate these new or refined annotations (Methods), which currently have very limited, and not rapid, incorporation into wide-spread annotation databases (104, 105). We are committed to keeping DRAM open to support community principles, with addition of new metabolisms fueled by community expertise. We call on any interested experts to join this endeavor and enable its continual development (link). Collectively, DRAM and DRAM-v deliver an infrastructure that enables rapid descriptions of microbial and viral contributions to ecosystem scale processes.

## AVAILABILITY

All DRAM source code is available at https://github.com/shafferm/DRAM under the GPL3 license. The DRAM user help is available at https://github.com/shafferm/DRAM/wiki. DRAM can also be installed via pip.

## ACCESSION NUMBERS

The *E. coli* genome was retrieved from KEGG. The set of 15 soil genomes were retrieved from NCBI. The Emerson *et al*. viral contigs were retrieved from GenBank, accession number QGNH00000000. The 44 gut metagenome samples in **Figure 4** were retrieved from HMP database. The single binned HMP gut metagenome sample used in **Figure 5** was retrieved from NCBI using accession number SRS019068, and the respective bins generated here deposited at NCBI. All accession numbers for MAGS and reads are detailed in **Supplementary File 4**.

## ACKNOWLEDGEMENTS

The following PIs (K.C.W., M.B.S., S.R.) and their respective affiliates are partially supported by funding from the National Sciences Foundation Division of Biological Infrastructure to build DRAM-v (Award# 1759874). DRAM was supported by the efforts of multiple individuals and grants within the Wrighton Laboratory. M.S., R.A.D, and K.C.W were partially funded by a National Institute of Health grant (HHS-NIH grant, Award# 007447-00002). P.L., A.B.N., K.C.W. were partially supported by an Early Career Award from the U.S. Department of Energy to K.C.W., Office of Science, Office of Biological and Environmental Research under Award Number DE-SC0018022. B.B.M was partially supported by an Early Career Award from the National Science Foundation to K.C.W., under Award Number 1750189. J.R.R is funded by the National Science Foundation grant (NRT-DESE, Award #1450032), support for A Trans-Disciplinary Graduate Training Program in Biosensing and Computational Biology at Colorado State University. The work conducted by the U.S. Department of Energy Joint Genome Institute is supported by the Office of Science of the U.S. Department of Energy under contract no. DE-AC02-05CH11231. The authors would also like to thank James Wainaina and Funing Tian for helpful discussion in the development of DRAM-v rules, as well as Tyson Claffey and Richard Wolfe for Colorado State server management. Computing resources for this work were also partially retained from The Ohio State University Unity cluster and the Ohio Supercomputer.

## AUTHOR INFORMATION

Michael Shaffer and Mikayla A. Borton contributed equally to this work.

## REFERENCES

1. Thompson, L.R., Sanders, J.G., McDonald, D., Amir, A., Ladau, J., Locey, K.J., Prill, R.J., Tripathi, A., Gibbons, S.M., Ackermann, G., et al. (2017) A communal catalogue reveals Earth’s multiscale microbial diversity. Nature, 551, 457–463.

2. Bolyen, E., Rideout, J.R., Dillon, M.R., Bokulich, N.A., Abnet, C.C., Al-Ghalith, G.A., Alexander, H., Alm, E.J., Arumugam, M., Asnicar, F., et al. (2019) Reproducible, interactive, scalable and extensible microbiome data science using QIIME 2. Nat. Biotechnol., 37, 852–857.

3. Wrighton, K.C., Thomas, B.C., Sharon, I., Miller, C.S., Castelle, C.J., VerBerkmoes, N.C., Wilkins, M.J., Hettich, R.L., Lipton, M.S., Williams, K.H., et al. (2012) Fermentation, hydrogen, and sulfur metabolism in multiple uncultivated bacterial phyla. Science (80-.)., 337, 1661–1665.

4. Sharon, I. and Banfield, J.F. (2013) Genomes from metagenomics. Science (80-.)., 342, 1057–1058.

5. Tyson, G.W., Chapman, J., Hugenholtz, P., Allen, E.E., Ram, R.J., Richardson, P.M., Solovyev, V. V, Rubin, E.M., Rokhsar, D.S. and Banfield, J.F. (2004) Community structure and metabolism through reconstruction of microbial genomes from the environment. Nature, 428, 37–43.

6. Angle, J.C., Morin, T.H., Solden, L.M., Narrowe, A.B., Smith, G.J., Borton, M.A., Rey-Sanchez, C., Daly, R.A., Mirfenderesgi, G., Hoyt, D.W., et al. (2017) Methanogenesis in oxygenated soils is a substantial fraction of wetland methane emissions. Nat. Commun., 8, 1567.

7. Woodcroft, B.J., Singleton, C.M., Boyd, J.A., Evans, P.N., Emerson, J.B., Zayed, A.A.F., Hoelzle, R.D., Lamberton, T.O., McCalley, C.K., Hodgkins, S.B., et al. (2018) Genome-centric view of carbon processing in thawing permafrost. Nature, 560, 49–54.

8. McCalley, C.K., Woodcroft, B.J., Hodgkins, S.B., Wehr, R.A., Kim, E.H., Mondav, R., Crill, P.M., Chanton, J.P., Rich, V.I., Tyson, G.W., et al. (2014) Methane dynamics regulated by microbial community response to permafrost thaw. Nature, 514, 478–481.

9. Roberts, A.B., Gu, X., Buffa, J.A., Hurd, A.G., Wang, Z., Zhu, W., Gupta, N., Skye, S.M., Cody, D.B., Levison, B.S., et al. (2018) Development of a gut microbe–targeted nonlethal therapeutic to inhibit thrombosis potential. Nat. Med., 24, 1407–1417.

10. Haiser, H.J., Gootenberg, D.B., Chatman, K., Sirasani, G., Balskus, E.P. and Turnbaugh, P.J. (2013) Predicting and manipulating cardiac drug inactivation by the human gut bacterium Eggerthella lenta. Science (80-.)., 341, 295–298.

11. Parks, D.H., Chuvochina, M., Waite, D.W., Rinke, C., Skarshewski, A., Chaumeil, P.A. and Hugenholtz, P. (2018) A standardized bacterial taxonomy based on genome phylogeny substantially revises the tree of life. Nat. Biotechnol., 36, 996.

12. Hug, L.A., Baker, B.J., Anantharaman, K., Brown, C.T., Probst, A.J., Castelle, C.J., Butterfield, C.N., Hernsdorf, A.W., Amano, Y., Ise, K., et al. (2016) A new view of the tree of life. Nat. Microbiol., 1, 16048.

13. Spang, A., Saw, J.H., Jørgensen, S.L., Zaremba-Niedzwiedzka, K., Martijn, J., Lind, A.E., Van Eijk, R., Schleper, C., Guy, L. and Ettema, T.J.G. (2015) Complex archaea that bridge the gap between prokaryotes and eukaryotes. Nature, 521, 173–179.

14. Emerson, J.B., Roux, S., Brum, J.R., Bolduc, B., Woodcroft, B.J., Jang, H. Bin, Singleton, C.M., Solden, L.M., Naas, A.E., Boyd, J.A., et al. (2018) Host-linked soil viral ecology along a permafrost thaw gradient. Nat. Microbiol., 3, 870.

15. Roux, S., Brum, J.R., Dutilh, B.E., Sunagawa, S., Duhaime, M.B., Loy, A., Poulos, B.T., Solonenko, N., Lara, E., Poulain, J., et al. (2016) Ecogenomics and potential biogeochemical impacts of globally abundant ocean viruses. Nature, 537, 689–693.

16. Nayfach, S., Shi, Z.J., Seshadri, R., Pollard, K.S. and Kyrpides, N.C. (2019) New insights from uncultivated genomes of the global human gut microbiome. Nature, 568, 505–510.

17. Pasolli, E., Asnicar, F., Manara, S., Zolfo, M., Karcher, N., Armanini, F., Beghini, F., Manghi, P., Tett, A., Ghensi, P., et al. (2019) Extensive Unexplored Human Microbiome Diversity Revealed by Over 150,000 Genomes from Metagenomes Spanning Age, Geography, and Lifestyle. Cell, 176, 649–662.

18. Almeida, A., Mitchell, A.L., Boland, M., Forster, S.C., Gloor, G.B., Tarkowska, A., Lawley, T.D. and Finn, R.D. (2019) A new genomic blueprint of the human gut microbiota. Nature, 568, 499.

19. Tully, B.J., Graham, E.D. and Heidelberg, J.F. (2018) The reconstruction of 2,631 draft metagenome-assembled genomes from the global oceans. Sci. Data, 5, 170203.

20. Stewart, R.D., Auffret, M.D., Warr, A., Walker, A.W., Roehe, R. and Watson, M. (2019) Compendium of 4,941 rumen metagenome-assembled genomes for rumen microbiome biology and enzyme discovery. Nat. Biotechnol., 37, 953–961.

21. Anantharaman, K., Brown, C.T., Hug, L.A., Sharon, I., Castelle, C.J., Probst, A.J., Thomas, B.C., Singh, A., Wilkins, M.J., Karaoz, U., et al. (2016) Thousands of microbial genomes shed light on interconnected biogeochemical processes in an aquifer system. Nat. Commun., 7, 13219.

22. Daly, R.A., Roux, S., Borton, M.A., Morgan, D.M., Johnston, M.D., Booker, A.E., Hoyt, D.W., Meulia, T., Wolfe, R.A., Hanson, A.J., et al. (2019) Viruses control dominant bacteria colonizing the terrestrial deep biosphere after hydraulic fracturing. Nat. Microbiol., 4, 352–361.

23. Al-Shayeb, B., Sachdeva, R., Chen, L.-X., Ward, F., Munk, P., Devoto, A., Castelle, C.J., Olm, M.R., Bouma-Gregson, K., Amano, Y., et al. (2020) Clades of huge phages from across Earth’s ecosystems. Nature, 578, 425–431.

24. Chaumeil, P.-A., Mussig, A.J., Hugenholtz, P. and Parks, D.H. (2018) GTDB-Tk: a toolkit to classify genomes with the Genome Taxonomy Database. Bioinformatics, 36, 1925–1927.

25. Bowers, R.M., Kyrpides, N.C., Stepanauskas, R., Harmon-Smith, M., Doud, D., Reddy, T.B.K., Schulz, F., Jarett, J., Rivers, A.R., Eloe-Fadrosh, E.A., et al. (2017) Minimum information about a single amplified genome (MISAG) and a metagenome-assembled genome (MIMAG) of bacteria and archaea. Nat. Biotechnol., 35, 725–731.

26. Jang, H. Bin, Bolduc, B., Zablocki, O., Kuhn, J.H., Roux, S., Adriaenssens, E.M., Brister, J.R., Kropinski, A.M., Krupovic, M., Lavigne, R., et al. (2019) Taxonomic assignment of uncultivated prokaryotic virus genomes is enabled by gene-sharing networks. Nat. Biotechnol., 37, 632–639.

27. Roux, S., Adriaenssens, E.M., Dutilh, B.E., Koonin, E. V., Kropinski, A.M., Krupovic, M., Kuhn, J.H., Lavigne, R., Brister, J.R., Varsani, A., et al. (2019) Minimum information about an uncultivated virus genome (MIUVIG). Nat. Biotechnol., 37, 29–37.

28. Heintz-Buschart, A., May, P., Laczny, C.C., Lebrun, L.A., Bellora, C., Krishna, A., Wampach, L., Schneider, J.G., Hogan, A., De Beaufort, C., et al. (2016) Integrated multi-omics of the human gut microbiome in a case study of familial type 1 diabetes. Nat. Microbiol., 2, 1–13.

29. Louca, S., Parfrey, L.W. and Doebeli, M. (2016) Decoupling function and taxonomy in the global ocean microbiome. Science (80-.)., 353, 1272–1277.

30. Seemann, T. (2014) Prokka: rapid prokaryotic genome annotation. Bioinformatics, 30, 2068–2069.

31. Tanizawa, Y., Fujisawa, T. and Nakamura, Y. (2018) DFAST: a flexible prokaryotic genome annotation pipeline for faster genome publication. Bioinformatics, 34, 1037–1039.

32. Dong, X. and Strous, M. (2019) An Integrated Pipeline for Annotation and Visualization of Metagenomic Contigs. Front. Genet., 10, 999.

33. Konwar, K.M., Hanson, N.W., Pagé, A.P. and Hallam, S.J. (2013) MetaPathways: a modular pipeline for constructing pathway/genome databases from environmental sequence information. BMC Bioinformatics, 14, 202.

34. Chen, I.-M.A., Chu, K., Palaniappan, K., Pillay, M., Ratner, A., Huang, J., Huntemann, M., Varghese, N., White, J.R., Seshadri, R., et al. (2019) IMG/M v. 5.0: an integrated data management and comparative analysis system for microbial genomes and microbiomes. Nucleic Acids Res., 47, D666–D677.

35. Moriya, Y., Itoh, M., Okuda, S., Yoshizawa, A.C. and Kanehisa, M. (2007) KAAS: an automatic genome annotation and pathway reconstruction server. Nucleic Acids Res., 35, W182–W185.

36. Aziz, R.K., Bartels, D., Best, A.A., DeJongh, M., Disz, T., Edwards, R.A., Formsma, K., Gerdes, S., Glass, E.M., Kubal, M., et al. (2008) The RAST Server: Rapid Annotations using Subsystems Technology. BMC Genomics, 9, 75.

37. Brum, J.R. and Sullivan, M.B. (2015) Rising to the challenge: accelerated pace of discovery transforms marine virology. Nat. Rev. Microbiol., 13, 147–159.

38. Hurwitz, B.L. and U’Ren, J.M. (2016) Viral metabolic reprogramming in marine ecosystems. Curr. Opin. Microbiol., 31, 161–168.

39. Enault, F., Briet, A., Bouteille, L., Roux, S., Sullivan, M.B. and Petit, M.-A. (2017) Phages rarely encode antibiotic resistance genes: a cautionary tale for virome analyses. ISME J., 11, 237–247.

40. Hyatt, D., Chen, G.L., LoCascio, P.F., Land, M.L., Larimer, F.W. and Hauser, L.J. (2010) Prodigal: Prokaryotic gene recognition and translation initiation site identification. BMC Bioinformatics, 11, 119.

41. Kanehisa, M., Furumichi, M., Tanabe, M., Sato, Y. and Morishima, K. (2017) KEGG: new perspectives on genomes, pathways, diseases and drugs. Nucleic Acids Res., 45, D353–D361.

42. Suzek, B.E., Wang, Y., Huang, H., McGarvey, P.B. and Wu, C.H. (2015) UniRef clusters: a comprehensive and scalable alternative for improving sequence similarity searches. Bioinformatics, 31, 926–932.

43. Rawlings, N.D., Barrett, A.J. and Bateman, A. (2010) MEROPS: the peptidase database. Nucleic Acids Res., 38, D227--D233.

44. Steinegger, M. and Söding, J. (2017) MMseqs2 enables sensitive protein sequence searching for the analysis of massive data sets. Nat. Biotechnol., 35, 1026–1028.

45. Daly, R.A., Borton, M.A., Wilkins, M.J., Hoyt, D.W., Kountz, D.J., Wolfe, R.A., Welch, S.A., Marcus, D.N., Trexler, R. V, MacRae, J.D., et al. (2016) Microbial metabolisms in a 2.5-km-deep ecosystem created by hydraulic fracturing in shales. Nat. Microbiol., 1, 16146.

46. El-Gebali, S., Mistry, J., Bateman, A., Eddy, S.R., Luciani, A., Potter, S.C., Qureshi, M., Richardson, L.J., Salazar, G.A., Smart, A., et al. (2019) The Pfam protein families database in 2019. Nucleic Acids Res., 47, D427–D432.

47. Finn, R.D., Clements, J. and Eddy, S.R. (2011) HMMER web server: interactive sequence similarity searching. Nucleic Acids Res., 39, W29--W37.

48. Zhang, H., Yohe, T., Huang, L., Entwistle, S., Wu, P., Yang, Z., Busk, P.K., Xu, Y. and Yin, Y. (2018) DbCAN2: A meta server for automated carbohydrate-active enzyme annotation. Nucleic Acids Res., 46, W95–W101.

49. Aramaki, T., Blanc-Mathieu, R., Endo, H., Ohkubo, K., Kanehisa, M., Goto, S. and Ogata, H. (2019) KofamKOALA: KEGG ortholog assignment based on profile HMM and adaptive score threshold. Bioinformatics, 36, 2251–2252.

50. Chan, P.P. and Lowe, T.M. (2019) tRNAscan-SE: Searching for tRNA genes in genomic sequences. In Methods in Molecular Biology. Humana Press Inc., Vol. 1962, pp. 1–14.

51. Parks, D.H., Imelfort, M., Skennerton, C.T., Hugenholtz, P. and Tyson, G.W. (2015) CheckM: assessing the quality of microbial genomes recovered from isolates, single cells, and metagenomes. Genome Res., 25, 1043–1055.

52. Vanderplas, J., Granger, B.E., Heer, J., Moritz, D., Wongsuphasawat, K., Satyanarayan, A., Lees, E., Timofeev, I., Welsh, B. and Sievert, S. (2018) Altair: Interactive Statistical Visualizations for Python. J. Open Source Softw., 3, 1057.

53. Gevers, D., Knight, R., Petrosino, J.F., Huang, K., McGuire, A.L., Birren, B.W., Nelson, K.E., White, O., Methé, B.A. and Huttenhower, C. (2012) The Human Microbiome Project: a community resource for the healthy human microbiome. PLoS Biol., 10, e1001377.

54. Bairoch, A. and Boeckmann, B. (1991) The SWISS-PROT protein sequence data bank. Nucleic Acids Res., 19, 2247.

55. Noguchi, H., Taniguchi, T. and Itoh, T. (2008) MetaGeneAnnotator: detecting species-specific patterns of ribosomal binding site for precise gene prediction in anonymous prokaryotic and phage genomes. DNA Res., 15, 387–396.

56. Huttenhower, C., Gevers, D., Knight, R., Abubucker, S., Badger, J.H., Chinwalla, A.T., Creasy, H.H., Earl, A.M., Fitzgerald, M.G., Fulton, R.S., et al. (2012) Structure, function and diversity of the healthy human microbiome. Nature, 486, 207–214.

57. Bushnell, B. (2018) BBTools. BBMap.

58. Peng, Y., Leung, H.C.M., Yiu, S.M. and Chin, F.Y.L. (2012) IDBA-UD: a de novo assembler for single-cell and metagenomic sequencing data with highly uneven depth. Bioinformatics, 28, 1420–1428.

59. Kang, D.D., Li, F., Kirton, E., Thomas, A., Egan, R., An, H. and Wang, Z. (2019) MetaBAT 2: An adaptive binning algorithm for robust and efficient genome reconstruction from metagenome assemblies. PeerJ, 2019, e7359.

60. Olm, M.R., Brown, C.T., Brooks, B. and Banfield, J.F. (2017) dRep: a tool for fast and accurate genomic comparisons that enables improved genome recovery from metagenomes through de-replication. ISME J., 11, 2864–2868.

61. Roux, S., Enault, F., Hurwitz, B.L. and Sullivan, M.B. (2015) VirSorter: mining viral signal from microbial genomic data. PeerJ, 3, e985.

62. Sullivan, M.B., Lindell, D., Lee, J.A., Thompson, L.R., Bielawski, J.P. and Chisholm, S.W. (2006) Prevalence and Evolution of Core Photosystem II Genes in Marine Cyanobacterial Viruses and Their Hosts. PLoS Biol., 4, e234.

63. Lindell, D., Jaffe, J.D., Johnson, Z.I., Church, G.M. and Chisholm, S.W. (2005) Photosynthesis genes in marine viruses yield proteins during host infection. Nature, 438, 86–89.

64. Frost, L.S., Leplae, R., Summers, A.O. and Toussaint, A. (2005) Mobile genetic elements: the agents of open source evolution. Nat. Rev. Microbiol., 3, 722–732.

65. Broecker, F. and Moelling, K. (2019) Evolution of immune systems from viruses and transposable elements. Front. Microbiol., 10, 51.

66. Gerlt, J.A., Bouvier, J.T., Davidson, D.B., Imker, H.J., Sadkhin, B., Slater, D.R. and Whalen, K.L. (2015) Enzyme function initiative-enzyme similarity tool (EFI-EST): A web tool for generating protein sequence similarity networks. Biochim. Biophys. Acta - Proteins Proteomics, 1854, 1019–1037.

67. Shannon, P., Markiel, A., Ozier, O., Baliga, N.S., Wang, J.T., Ramage, D., Amin, N., Schwikowski, B. and Ideker, T. (2003) Cytoscape: a software environment for integrated models of biomolecular interaction networks. Genome Res., 13, 2498–504.

68. Bland, C., Ramsey, T.L., Sabree, F., Lowe, M., Brown, K., Kyrpides, N.C. and Hugenholtz, P. (2007) CRISPR Recognition Tool (CRT): a tool for automatic detection of clustered regularly interspaced palindromic repeats. BMC Bioinformatics, 8, 209.

69. UniProt: the universal protein knowledgebase (2017) Nucleic Acids Res., 45, D158--D169.

70. Galperin, M.Y. and Koonin, E. V (2004) ‘Conserved hypothetical’proteins: prioritization of targets for experimental study. Nucleic Acids Res., 32, 5452–5463.

71. Smith, G.J., Angle, J.C., Solden, L.M., Borton, M.A., Morin, T.H., Daly, R.A., Johnston, M.D., Stefanik, K.C., Wolfe, R., Gil, B., et al. (2018) Members of the Genus Methylobacter Are Inferred To Account for the Majority of Aerobic Methane Oxidation in Oxic Soils from a Freshwater Wetland. MBio, 9, e00815–18.

72. Tavormina, P.L., Orphan, V.J., Kalyuzhnaya, M.G., Jetten, M.S.M. and Klotz, M.G. (2011) A novel family of functional operons encoding methane/ammonia monooxygenase-related proteins in gammaproteobacterial methanotrophs. Environ. Microbiol. Rep., 3, 91–100.

73. Kelley, L.A., Mezulis, S., Yates, C.M., Wass, M.N. and Sternberg, M.J.E. (2015) The Phyre2 web portal for protein modeling, prediction and analysis. Nat. Protoc., 10, 845–858.

74. Wrighton, K.C., Castelle, C.J., Varaljay, V.A., Satagopan, S., Brown, C.T., Wilkins, M.J., Thomas, B.C., Sharon, I., Williams, K.H., Tabita, F.R., et al. (2016) RubisCO of a nucleoside pathway known from Archaea is found in diverse uncultivated phyla in bacteria. ISME J., 10, 2702.

75. Hobbs, E.T., Pereira, T., O’Neill, P.K. and Erill, I. (2016) A Bayesian inference method for the analysis of transcriptional regulatory networks in metagenomic data. Algorithms Mol. Biol., 11, 19.

76. Imachi, H., Nobu, M.K., Nakahara, N., Morono, Y., Ogawara, M., Takaki, Y., Takano, Y., Uematsu, K., Ikuta, T., Ito, M., et al. (2020) Isolation of an archaeon at the prokaryote–eukaryote interface. Nature, 577, 519–525.

77. Vignais, P.M. and Billoud, B. (2007) Occurrence, classification, and biological function of hydrogenases: An overview. Chem. Rev., 107, 4206–4272.

78. Mehta, R.S., Abu-Ali, G.S., Drew, D.A., Lloyd-Price, J., Subramanian, A., Lochhead, P., Joshi, A.D., Ivey, K.L., Khalili, H., Brown, G.T., et al. (2018) Stability of the human faecal microbiome in a cohort of adult men. Nat. Microbiol., 3, 347–355.

79. Johnson, A.J., Vangay, P., Al-Ghalith, G.A., Hillmann, B.M., Ward, T.L., Shields-Cutler, R.R., Kim, A.D., Shmagel, A.K., Syed, A.N., Students, P.M.C., et al. (2019) Daily sampling reveals personalized diet-microbiome associations in humans. Cell Host Microbe, 25, 789–802.

80. Baker, D., Norris, K.H. and Li, B.W. (1979) Food Fiber Analysis: Advances in Methodology in Dietary Fibers: Chemistry and Nutrition. GW Inglett and SL Falkehog.

81. Maxwell, E.G., Belshaw, N.J., Waldron, K.W. and Morris, V.J. (2012) Pectin--an emerging new bioactive food polysaccharide. Trends Food Sci. Technol., 24, 64–73.

82. Stefler, D. and Bobak, M. (2015) Does the consumption of fruits and vegetables differ between Eastern and Western European populations? Systematic review of cross-national studies. Arch. Public Heal., 73, 29.

83. Zeevi, D., Korem, T., Zmora, N., Israeli, D., Rothschild, D., Weinberger, A., Ben-Yacov, O., Lador, D., Avnit-Sagi, T., Lotan-Pompan, M., et al. (2015) Personalized Nutrition by Prediction of Glycemic Responses. Cell, 163, 1079–1094.

84. Korem, T., Zeevi, D., Zmora, N., Weissbrod, O., Bar, N., Lotan-Pompan, M., Avnit-Sagi, T., Kosower, N., Malka, G., Rein, M., et al. (2017) Bread Affects Clinical Parameters and Induces Gut Microbiome-Associated Personal Glycemic Responses. Cell Metab., 25, 1243–1253.e5.

85. Lozupone, C.A., Stombaugh, J.I., Gordon, J.I., Jansson, J.K. and Knight, R. (2012) Diversity, stability and resilience of the human gut microbiota. Nature, 489, 220–230.

86. Amaretti, A., Gozzoli, C., Simone, M., Raimondi, S., Righini, L., Pérez-Brocal, V., García-López, R., Moya, A. and Rossi, M. (2019) Profiling of Protein Degraders in Cultures of Human Gut Microbiota. Front. Microbiol., 10, 2614.

87. Chambers, E.S., Preston, T., Frost, G. and Morrison, D.J. (2018) Role of Gut Microbiota-Generated Short-Chain Fatty Acids in Metabolic and Cardiovascular Health. Curr. Nutr. Rep., 7, 198–206.

88. Tran, H.T., Barnich, N. and Mizoguchi, E. (2011) Potential role of chitinases and chitin-binding proteins in host-microbial interactions during the development of intestinal inflammation. Histol. Histopathol., 26, 1453.

89. Filpa, V., Moro, E., Protasoni, M., Crema, F., Frigo, G. and Giaroni, C. (2016) Role of glutamatergic neurotransmission in the enteric nervous system and brain-gut axis in health and disease. Neuropharmacology, 111, 14–33.

90. Tang, W.H.W., Li, D.Y. and Hazen, S.L. (2019) Dietary metabolism, the gut microbiome, and heart failure. Nat. Rev. Cardiol., 16, 137–154.

91. Turnbaugh, P.J., Ley, R.E., Mahowald, M.A., Magrini, V., Mardis, E.R. and Gordon, J.I. (2006) An obesity-associated gut microbiome with increased capacity for energy harvest. Nature, 444, 1027.

92. Wu, J., Sabag-Daigle, A., Borton, M.A., Kop, L.F.M., Szkoda, B.E., Kaiser, B.L.D., Lindemann, S.R., Renslow, R.S., Wei, S., Nicora, C.D., et al. (2018) Salmonella-mediated inflammation eliminates competitors for fructose-asparagine in the gut. Infect. Immun., 86, e00945–17.

93. Solden, L.M., Naas, A.E., Roux, S., Daly, R.A., Collins, W.B., Nicora, C.D., Purvine, S.O., Hoyt, D.W., Schückel, J., Jørgensen, B., et al. (2018) Interspecies cross-feeding orchestrates carbon degradation in the rumen ecosystem. Nat. Microbiol., 3, 1274.

94. Breitbart, M., Thompson, L., Suttle, C. and Sullivan, M. (2007) Exploring the Vast Diversity of Marine Viruses. Oceanography, 20, 135–139.

95. Mizuno, C.M., Guyomar, C., Roux, S., Lavigne, R., Rodriguez-Valera, F., Sullivan, M.B., Gillet, R., Forterre, P. and Krupovic, M. (2019) Numerous cultivated and uncultivated viruses encode ribosomal proteins. Nat. Commun., 10, 752.

96. Garneau, J.R., Depardieu, F., Fortier, L.-C., Bikard, D. and Monot, M. (2017) PhageTerm: a tool for fast and accurate determination of phage termini and packaging mechanism using next-generation sequencing data. Sci. Rep., 7, 1–10.

97. Reese, A.T., Pereira, F.C., Schintlmeister, A., Berry, D., Wagner, M., Hale, L.P., Wu, A., Jiang, S., Durand, H.K., Zhou, X., et al. (2018) Microbial nitrogen limitation in the mammalian large intestine. Nat. Microbiol., 3, 1441–1450.

98. Zengler, K. and Zaramela, L.S. (2018) The social network of microorganisms - How auxotrophies shape complex communities. Nat. Rev. Microbiol., 16, 383–390.

99. Holeček, M. (2018) Branched-chain amino acids in health and disease: metabolism, alterations in blood plasma, and as supplements. Nutr. Metab. (Lond)., 15, 33.

100. Dwight, S.S., Harris, M.A., Dolinski, K., Ball, C.A., Binkley, G., Christie, K.R., Fisk, D.G., Issel-Tarver, L., Schroeder, M., Sherlock, G., et al. (2002) Saccharomyces Genome Database (SGD) provides secondary gene annotation using the Gene Ontology (GO). Nucleic Acids Res., 30, 69–72.

101. Trifinopoulos, J., Nguyen, L.-T., von Haeseler, A. and Minh, B.Q. (2016) W-IQ-TREE: a fast online phylogenetic tool for maximum likelihood analysis. Nucleic Acids Res., 44, W232--W235.

102. Yu, N.Y., Wagner, J.R., Laird, M.R., Melli, G., Rey, S., Lo, R., Dao, P., Sahinalp, S.C., Ester, M., Foster, L.J., et al. (2010) PSORTb 3.0: improved protein subcellular localization prediction with refined localization subcategories and predictive capabilities for all prokaryotes. Bioinformatics, 26, 1608–1615.

103. Robinson, J.T., Thorvaldsdóttir, H., Winckler, W., Guttman, M., Lander, E.S., Getz, G. and Mesirov, J.P. (2011) Integrative genomics viewer. Nat. Biotechnol., 29, 24–26.

104. Ticak, T., Kountz, D.J., Girosky, K.E., Krzycki, J.A. and Ferguson, D.J. (2014) A nonpyrrolysine member of the widely distributed trimethylamine methyltransferase family is a glycine betaine methyltransferase. Proc. Natl. Acad. Sci., 111, E4668--E4676.

105. Craciun, S. and Balskus, E.P. (2012) Microbial conversion of choline to trimethylamine requires a glycyl radical enzyme. Proc. Natl. Acad. Sci., 109, 21307–21312.

