## Supplementary Information for "DRAM for distilling microbial metabolism to automate the curation of microbiome function"

**Online Resources:**

DRAM and DRAM-v source code and documentation:
https://github.com/shafferm/DRAM

DRAM and DRAM-v user guide:
https://github.com/shafferm/DRAM/wiki

**Table of Contents**

Online Resources ........................................ **1**

Table of Contents ........................................ **2**

Supplementary Methods

*Detailed DRAM and DRAM-v output description* ........................................ **3-5**

*Data sets used in this manuscript to evaluate DRAM performance* ........................................ **5-6**

*Benchmarking and performance analysis* ........................................ **6-10**

*DRAM pathways and enzyme modularity completion* ........................................ **10-11**

*Interpreting results from DRAM and DRAM-v* ........................................ **11-15**

Supplementary Discussion

*Comparison of DRAM features to other tools*........................................**16-17**

*Measuring Precision and Recall of DRAM annotations using KEGG Genes and E. coli* .......**17-18**

*Dechloromonas aromatica RCB full genome description* ........................................ **18**

*Integration of DRAM and DRAM-v to begin to infer virocell metabolism*........................................ **18-20**

Supplementary Tables

*Comparison of features across several annotators* ........................................ **21**

Supplementary Figures ........................................ **22-34**

Supplementary Files Descriptions ........................................ **35-36**

Supplementary References ........................................ **37-41**

### SUPPLEMENTARY METHODS

#### ***Detailed DRAM and DRAM-v output file description***

Below we outline the multiple output files for DRAM and DRAM-v, as shown in **Figure 1**. The content of each output file is detailed below:

DRAM microbial genome output files

(1) *Raw* (obtained by running DRAM.py annotate)

- Tab separated file (.tsv) with all the annotations from Pfam (1), KEGG (2), UniProt90 (3), VOGDB (http://vogdb.org/), dbCAN (4), and MEROPS (5) databases for all genes in all input genomes
- GenBank (.gbk) files for each genome
- General feature format (GFF) (.gff) file of all annotations across genomes
- FASTA format file (.fna) of each open reading frame nucleotide sequence and best ranked annotation (see Annotation ranks section)
- FASTA format file (.faa) of each translated open reading frame amino acid sequence and best ranked annotation KEGG (2) annotation
- FASTA format file (.fasta) of all entries from all input FASTA files with names matching those in the GFF file

(2) *Distillate* (obtained by running DRAM.py distill)

- Tab separated file (.tsv) with genome statistics for all input genomes including all mandatory genome quality statistics required by recently defined MIMAG standards (6)
- Microsoft Excel spreadsheet (.xlsx) with metabolism summary of all input genomes, which gives gene counts of functional and structural genes across a wide variety of metabolisms

(3) *Product* (obtained by running DRAM.py distill)

- HTML (.html) file containing an interactive heatmap showing coverage of pathways, the coverage of electron transport chain components, and the presence of selected metabolic functions
- Tab separated file (.tsv) with data from the html heatmap.

DRAM-v for viral contigs and genomes output

(1) *Raw* (obtained by running DRAM-v.py annotate)

- Tab separated file (.tsv) with all the annotations from PFAM (1), KEGG (2), Uniref90 (3), dbCAN (4), MEROPS (5), VOGDB (http://vogdb.org/), and the viral subset of RefSeq (7) for all genes
- Folder of GenBank files with annotations for each viral contig
- General feature format (.gff) file of all annotations across viral contigs
- FASTA format file (.fna) of each open reading frame nucleotide sequence and best ranked annotation
- FASTA format file (.faa) of each translated open reading frame amino acid sequence and best ranked annotation

(2) *Distillate* (obtained by running DRAM-v.py distill)

- Tab separated file (.tsv) with viral contig statistics for all input viral contigs including some statistics required by recently defined MIUVIG standards (8)
- Tab separated file (.tsv) of auxiliary metabolic genes (AMG) summary from all input viral contigs, which lists putative AMGs with annotation, auxiliary scores, and other flags outlined in main text **Figure 6ab.**

(3) *Product* (obtained by running DRAM-v.py distill)

- HTML (.html) file containing an interactive heatmap showing all viruses with a putative AMG and the AMG metabolism category, with number of AMGs on each contig noted

- Tab separated file (.tsv) with corresponding AMGs from the html heatmap.

In summary, DRAM provides the user with several outputs, including many formats that allow the user to summarize genome metabolic properties and visualize this data quickly. Additionally, we note that DRAM is just the first step in functional gene annotation, thus we provide users with output files to ensure interoperability with other subsequent programs (e.g. GenBank, GFF, or tab delimited). This is different from many other annotation systems which stop at the *Raw* annotation step (Prokka (9), DFAST (10)), or are not designed to scale synthesized pathway analyses across multiple (to thousands) genomes (MetaErg (11)) (**Supplementary Table 1**). We acknowledge that every tool has strengths and limitations, and an optimal pipeline would likely use multiple tools simultaneously to maximize annotation understanding.

***Data sets used in this manuscript to evaluate DRAM performance***

All data used to evaluate DRAM performance, validate DRAM metabolic characterization, and showcase the capabilities of DRAM were published previously. The reasoning for the selection of each file is described below:

(1) *E. coli* K-12 MG1655 was used due to its complete annotations (NCBI Reference Sequence: NC_000913.3).

(2) An *in silico* constructed, representative soil microbial community composed of 15 isolate genomes and MAGs that capture the breadth of biogeochemical processes (e.g. methanotrophy, methanogenesis, denitrification, nitrification, photosynthesis, aerobic respiration, sulfur reduction and oxidation, nitrogen fixation and fermentation) encoded in phylogenetically distinct organisms. The metabolism represented by and the reasoning for choosing each isolate genome or MAG is detailed in **Supplementary File 4**.

(3) Human Microbiome Project (HMP) (12) sample SRS019068 was chosen to highlight the feasibility of DRAM to analyze more complex, authentic microbiome samples. We selected this

particular human fecal HMP sample because it was the largest available fecal metagenome in the HMP database (29 Gbp).

(4) To recreate the stability of function figure **(Figure 4a)** from the Human Microbiome Consortium paper published in 2012 (13), we randomly selected a subset of 44 metagenomes from the previously published analysis. For this subset, we assembled the samples separately using IDBA-UD (14) with default parameters and subsequently annotated with DRAM. We selected this approach to demonstrate how substrate-specific annotations enabled by DRAM offer new insights compared to the COG categories used historically. Accession numbers for the HMP metagenomes are listed in **Supplementary file 4**.

(5) To validate AMGs assigned by DRAM-v, we used a soil viral file that was previously published by our team (15) and included 14 putative AMGs that were called manually. DRAM-v recovered all 14 of the manually-called AMGs and an additional 453 high and medium quality AMGs.

(6) To demonstrate the scalability of DRAM to thousands of MAGs, we used DRAM to annotate one of the largest, published MAG collections from a single ecosystem (doi: 10.1038/ncomms13219, (16)). For these 2,535 MAGs, we provide the DRAM annotations (Raw, Distillate, and Product) outputs here: https://zenodo.org/record/3777237).

***Benchmarking and performance analysis***

Performance and benchmarking analysis for DRAM was done using 3 separate datasets (datasets 1-3 above). The number of (i) DRAM databases and their sequence entries, (ii) annotations, (iii) hypothetical genes, as well as the memory use and run time were compared to Prokka (9), DFAST (10), and MetaErg (11) annotators. All performance metrics for each annotator are detailed in **Supplementary File 4**. Here we define the following metrics across the 4 annotation tools, including DRAM:

- Annotation- we define a positive annotation of a given gene as:

  - Prokka (9): at least one annotation of the gene was not given as hypothetical", "uncharacterized" or "domain of unknown function"
  - DFAST (10): at least one annotation of the gene was not given as hypothetical", "uncharacterized" or "domain of unknown function"
  - MetaErg (11): at least one annotation of the gene from Swiss-Prot (17), TIGRFAM (18) or Pfam (1) was not hypothetical", "uncharacterized" or "domain of unknown function"
  - DRAM: at least one annotation of the gene from (bit score >60) KEGG (2), UniRef90 (3), MEROPs (5), Pfam (1) or dbCAN was not to a "hypothetical", "uncharacterized" or "domain of unknown function" gene
- Hypothetical- we define a hypothetical annotation of a given gene:
  - Prokka* (9): none (see Methods)
  - DFAST* (10): none (see Methods)
  - MetaErg (11): hits for a gene lacked defined annotation, and at least one of the annotations (from Swiss-Prot (17), TIGRFAM (18) and Pfam (1) databases) included terms "hypothetical", "uncharacterized" or "domain of unknown function".
  - DRAM: hits for a gene lacked defined annotation, and at least one of the annotations (from KEGG (2), UniRef90 (3), MEROPs (5), Pfam (1) and dbCAN2 (4)) included terms "hypothetical", "uncharacterized" or "domain of unknown function" terms.

Unannotated- we define an unannotated gene as:

  - Prokka* (9): annotation was listed as hypothetical
  - DFAST* (10): annotation was listed as hypothetical
  - MetaErg (11): no annotation was assigned from Swiss-Prot (17), TIGRFAM (18) or Pfam (1) databases

    - DRAM: no annotation was assigned from KEGG (2), UniRef90, MEROPs (5), Pfam (1) or dbCAN2 (4) databases
- Memory- we compare maximum memory in gigabytes needed to annotate each dataset across 4 annotation tools using default parameters.
- Run time- We compare total runtime in minutes to annotate each dataset across 4 annotation tools using default parameters.

* Prokka (9) and DFAST (10) remove hypothetical genes from their databases and subsequently call all genes without hits as hypothetical, when they could in fact be unannotated or hypothetical and there is no way for the user to know this. As such, in these DRAM performance analyses we considered DFAST and Prokka hypothetical labels as unannotated, as it was not possible to discern the difference between hypothetical and unannotated genes.

Performance evaluation of DRAM compared to other annotation tools using the *E. coli* strain K-12 MG1655 isolate genome showed that DRAM performed the best as measured by number of annotated and unannotated genes (when annotating with UniRef90 (3)): 4,158 annotated genes, 154 hypothetical genes, and only 4 genes left unannotated. MetaErg (11) performed second best with 4,001 annotated genes, 167 hypothetical genes, and 27 unannotated genes. Prokka (9) and DFAST (10) both performed similarly with 3,518 and 3,557 annotated and 787 and 749 unannotated genes, respectively. Time and memory usage metrics for annotation of this single *E. coli* genome showed that DFAST (10) was fastest (average 2.1 minutes), with Prokka (9) (average 8.4 minutes) and DRAM (average 12.7 minutes) close behind. MetaErg (11) was slowest at an average of 22.2 minutes (**Supplementary Figure 3b**).

The *E. coli* K-12 MG1655 genome was also used to evaluate the precision and recall of DRAM using default parameters. Specifically, we annotated this genome with DRAM using the full KEGG Genes database (2) and then again using three subsets of the KEGG Genes database (2) to test for KO annotation recovery. The subsets used were 1) KEGG Genes (2) with the *E. coli* K-12 MG1655 genes removed, 2) KEGG Genes (2) with all genes from all *E. coli* genomes removed and 3) KEGG Genes (2) with all genes from all Escherichia genomes removed. To evaluate performance annotations from DRAM were compared to annotations from the KEGG *E. coli* K-12 MG1655 genome. An annotation was considered a true positive if DRAM called and annotated the gene with the exact same set of KO identifiers as in the KEGG genome. An annotation was considered a false positive if the gene was called by DRAM and annotated with any set of KOs different from those in the KEGG genome. An annotation was considered a false negative if a gene was not called by DRAM but was in the KEGG genome or the gene was called by DRAM and not annotated with a KO identifier. The true positive, false positive and false negative rates were calculated for each KEGG Genes database (2) subset and used to calculate precision, recall and F1 score.

For the set of 15 soil genomes, performance results were similar to the *E. coli* genome, but allowed us to measure performance across taxonomically diverse members (**Supplementary File 4**). Across the entire set, Prokka (9) and DFAST (10) again performed similarly (25,466 and 31,258 total annotated genes, respectively), while MetaErg (11) did better with 42,478 hypothetical genes. Notably, DRAM annotated the most genes (45,662). More specifically, DRAM excelled compared to these annotators on the annotation recovery from uncultivated lineages (e.g. Candidate Phyla Radiation). This was exemplified by the Parcubacteria MAG, for which DRAM was able to annotate 826 genes, as well as recovered 361 hypothetical genes and 7 unannotated genes. Prokka (9) recovered 726 unannotated genes while DFAST (10) and MetaErg (11) left 755 and 342 genes, respectively. In other words, DRAM annotated approximately 2.7 (DFAST (10)), 2.5 (Prokka (9)) and 1.5 (MetaErg (11)) times more genes than the other tools. Using these multiple, disparate databases, DRAM had the second longest run time on average across five runs at requiring 209.9 minutes, with Prokka (9) and DFAST (10) at 27.0 and 32.6 minutes, respectively. MetaErg (11) took the longest with a runtime of 264.6 minutes on average.

Performance results from 76 MAGs generated from one Human Microbiome Project (12) metagenome (SRS019068) were similar to the *in silico* soil community, with DRAM having more combined annotated and hypothetical genes and less unannotated genes relative to other annotators (**Supplementary Figure 3c**). DRAM and MetaErg (11) had similar (595.9 and 563.7 minutes, respectively), yet slower runtimes compared to Prokka (9) (70.9 minutes) and DFAST (10) (78 minutes).

DRAM's maximum memory usage is unrelated to the amount of input data. Across all files evaluated here, DRAM had a maximum memory usage of 99GB, which is more than any other annotator. Specifically, Prokka (9), DFAST (10), and MetaErg (11) have a maximum memory usages of 1.0GB, 1.9GB, and 9.7GB (respectively).

DRAM's ability to recover more annotations and fewer unannotated genes due to expanded and disparate databases comes at the expense of runtime and memory. DRAM has at least 21- to 104-million more database entries, when annotating against UniRef90 (3), than Prokka (9), DFAST (10), and MetaErg (11) (**Supplementary File 4**). When comparing the single database that all annotation tools shared, Uniprot (19), DRAM has the most entries through UniRef90 (3). Other annotators have proprietarily culled this database resulting in 132-3,412 fold reduction in the number of entries from Uniprot (19) (**Figure 2cd**). Moreover, the annotator with the next largest set of database entries, MetaErg (11), uses GenomeDB which is based on the set of genomes from the Genome Taxonomy Database (20) annotated with NCBI RefSeq which further limits the tools ability to annotate genomes which are not well characterized. To overcome the time and memory cost of using the full UniRef90 (3) set DRAM does not annotate against UniRef90 (3)by default but this option can be selecting using the --use_uniref flag. By default, UniRef90 (3) annotations are skipped resulting in less annotations in the *Raw* output, but this will currently not impact the *Distillate* or *Product* reports.

***DRAM pathways and enzyme modularity completion***

DRAM surpasses other annotation tools by turning a list of genes in a genome, or even more unwieldly, genes across multiple genomes, into an inventory of genes known to catalyze biogeochemical reactions. This synthesized analysis is showcased in the DRAM *Product* (**Supplementary File 3**, **Figure 3**), which summarizes metabolic information in 3 ways: (1) pathway completion (e.g. glycolysis), (2) multi-subunit enzyme subunit completion (e.g. NADH dehydrogenase), and (3) presence/absence of process-defining functional gene(s) (e.g. xyloglucan degradation) (**Supplementary Figure 2**). Pathway completion considers unique routes for processes that start and end with the same compounds, while also accounting for different genes (shown by black circle outlines in **Supplementary Figure 2**) at each step.

In the *Product* the completion electron train complexes (ETC) are shown, as this completion information is critical for inferring differences in respiration machinery, as well as in annotating obligate fermenters (often absence of this machinery). with the completion of each complex reported. Completion is based on the number of subunits present for a given enzyme. This allows the user to discern which complexes are present in the ETC quickly, as well as interpret possible misannotations. The latter is demonstrated in **Supplementary Figure 2** by NADH dehydrogenase. We can infer that an organism with the full NADH dehydrogenase complex (14 out of 14 genes present, or 11/11) has the first complex in the electron transport chain, but an organism with partial completion, should be further investigated. For instance, the partial completion (3 genes) of the multi-subunit NADH dehydrogenase shown in **Supplementary Figure 2** is not due to a complete complex I, but rather the presence of NiFe hydrogenase common in obligate fermenters (21, 22). These hydrogenases are further annotated in detail by their type and function in the *Distillate* (**Supplementary File 2**).

***Interpreting results from DRAM and DRAM-v***

**I) Genome quality metrics and reporting**

The genome_stats.tsv is a part of the DRAM *Distillate* that compiles the statistics necessary for publishing bacterial, archaeal, and viral genomes (6, 8), making this time-consuming step of genome reporting automated. This file contains completeness, contamination, taxonomy, rRNAs present, tRNA count, number of scaffolds, and draft genome quality for each input genome. Based on these metrics, DRAM assigns quality scores of high medium and low according to MIMAG standards (6). Similarly vMAG_stats.tsv in the DRAM-v Distillate contains statistics for publishing vMAGs and the VIRSorter (20) category for each vMAG, if the contig is circular (closed genome), if the contig is a predicted prophage, gene count on the contig, the number of strand switches, the potential AMG count, if a transposase present on the contig, if this is a possible non-viral contig as well as the number of viral hypothetical genes, viral genes with unknown function, viral replication genes, viral structure genes, viral genes with host benefits, and viral genes with viral benefits according to VOGDB.

DRAM can accept MAG quality control metrics and taxonomy information from any outside source in the form of tab separated files with MAG names as the first column. DRAM looks for columns called 'Completeness' and 'Contamination' for quality information (this allows DRAM to accept results files from checkM (23) with no modification). Similarly for taxonomy DRAM looks for a column called 'classification' for taxonomy assignment (this allows DRAM to accept results files from GTDB-tk (20) with no modification).

All other genome metrics are calculated within DRAM including rRNAs present using barrnap (https://github.com/tseemann/barrnap), tRNAs using tRNA-scan (24), and number of scaffolds. Beyond genome quality statistics, DRAM also exports GenBank files for each genome and a GFF file with all annotated genomes.

### II) General information on interpreting DRAM output

DRAM and DRAM-v are powerful first steps in profiling functionally important genes in assembled genomic data including a genome, set of genomes, or an unbinned, but assembled metagenome (scaffold list). As is the case with all homology based annotations, we recommend users confirm these initial DRAM annotations using a combination of methods not limited to: protein phylogenetic analyses, localization analyses (PSORTb (25)), confirmation of active sites, protein structure homology-modelling, and sequence similarity networks.

### III) Multi-heme c-type cytochromes

Iron-reducing and oxidizing microorganisms, among others, use multi-heme c-type cytochromes (MHCs) as electron transfer between the cell and mineral phase (21, 26). DRAM detects the presence of MHCs by reporting the number of CXXCH motifs present in each gene. This information can be found in the annotations.tsv output files. We note, this is the first step in identifying putative MHCs, thus to further validate gene as playing a role in mineral reduction, users should confirm the annotation (e.g. nitrate reductases also have multiple hemes in a gene), upload the gene sequence into PSORTb (25) to obtain cellular location information, and perform sequence similarity network analysis relative to known MHCs to better identify genes involved in extracellular metal respiration.

**IV) Methanogenesis**

The full methanogenesis pathway and specific genes crucial for determining substrate type are outlined in the DRAM *Distillate* and *Product*. For the pathway completion, DRAM accounts for the minimum 8 steps required for methanogenesis. Note, this does not include all the genes for input substrates (e.g. TMA, acetate), as it is possible that an organism uses a single specific substrate. The first step for inferring a methanogen is the presence of *mcr* gene, a functional gene involved in methane generation (included in *Product*). After that, we inventory some genes used by methanogens for substrate incorporation into methanogenesis (however these genes do not define a methanogen, as many homologs to these genes are also used by non-methanogens). The presence of the substrate specific genes are noted in the presence absence portion of the *Product* heatmap. For methylamine utilization, users should confirm the presence of Pyrrolysine in the methyltransferases, as non-Pyrrolysine containing homologs have been shown to demethylate other substrates (27, 28). For example, non-Pyrrolysine containing homologs of trimethylamine methyltransferase encoded in the genome *Desulfitobacterium hafninese* have been shown to demethylate glycine betaine instead of trimethylamine (27).

**IV) CAZymes**

The creation of the CAZy database and associated tools that utilize the resource (such as dbCAN (4)) has allowed users to identify carbohydrate active enzymes (CAZymes) in isolates, environmental metagenomes, and MAGs (15, 29–31). Here, we manually curated the CAZyme database (via dbCAN2 (4)) to provide the user with substrate-resolved metabolic potential, rather than just CAZy identifiers (e.g. glycoside hydrolase, GH#) that must be searched in the CAZy database to gain metabolic context. We provide broad level substrate classes (e.g. pectin) that allow a user to assign GHs into substrate classes, highlighting the key carbohydrate functions across genomes. In order to be considered able to degrade a specific substrate class in the *Product* heatmap, a genome must have BOTH a backbone or endo-cleaving CAZyme for the specific substrate AND an oligo or exo-cleaving CAZyme. However, as DRAM provides all levels of information, genomes that have either a backbone or endo-cleaving GH for the specific substrate OR an oligo or exo-cleaving glycoside hydrolase will be noted in the *Distillate* output (allowing for user identification of other bacteria to potentially cross-feed on specific substrates).

DRAM has eliminated the manual and time-consuming step of classifying CAZymes by (1) pulling the complete activity of each CAZyme family directly into the *Raw* and *Distillate* DRAM output so all annotations are in a single place, (2) accounting for the fact that some CAZymes can operate on more than one substrate, and (3) providing cleavage information to reduce false positives in carbon utilization (**Supplementary File 4**). Specific examples for each curation step are given below:

(1) For GH35, the DRAM *Raw* and *Distillate* output give you the annotation as beta-galactosidase (EC 3.2.1.23); exo-beta-glucosaminidase (EC 3.2.1.165); exo-beta-1,4-galactanase (EC 3.2.1.-); beta-1,3-galactosidase (EC 3.2.1.-) and oligo cleavage of beta-galactan.

(2) DRAM *Product* accounts for CAZymes that operate on more than one substrate: for example, GH5 has the potential to degrade xylans, mannans, beta-glucans, xyloglucans, amorphous cellulose and oligosaccharides. Alternatively, DRAM accounts for multiple CAZymes that can operate on a single substrate: phenolics can be degraded by AA1, AA2, or AA4 CAZyme families. Moreover, for phenolics, we have also included KO numbers to increase annotation coverage, as CAZy AA families are biased towards fungal enzymes (30): K05909, K05810, K00422.

(3) DRAM only annotates substrate utilization based on the presence of endo- and exo-cleavage information on both poly- and oligosaccharides. This interpretation is conservative, and was designed to reduce false positive annotations. For example, to be positive for pectin degradation in the DRAM *Product* heatmap, you have to have one of the back-bone cleaving CAZymes (GH28, PL1, PL2, PL9, PL10, PL11, PL22, PL26, PL3, PL4, PL9) and an oligo-cleaving CAZyme (GH78, GH106, GH138, GH139, GH143, CE8, GH105, GH4, CE12, GH2).

Moreover, we have not considered families that have very rare instances of an activity (or dual function) compared to a dominating function for defined substrate utilization in the *Product*. For example, there are very few cases of GH5, GH16 and GH30 displaying activity on galactans, thus these enzymes have been excluded from the list for these substrates. However, in the *Distillate* all CAZyme information and families in the database are reported.

**V) Peptidases**

DRAM profiles the putative peptidases and peptidase inhibitors in each input genome using information from the MEROPs database (5). Specifically, DRAM reports the peptidase catalytic type, family information, and specific protein or peptide interactions. This information is summarized per genome in the distillate output of DRAM, enabling the user to quantify the number and type of peptidase in a streamlined manner. For DRAM-v specifically, peptidases that have been commonly associated with viruses according to the MEROPs database (5) are not considered by DRAM to be AMGs and are thus removed from the putative AMG summary in DRAM-v.

**VI) AMGs**

DRAM-v profiles putative AMGs found within viral genomes. As shown in main text **Figure 6**, DRAM assigns an auxiliary score to viral genes, denoting how likely a putative metabolic gene is of viral origin (ranked from 1 on scale to 5). DRAM-v also flags possible metabolic genes that are experimentally verified or reported AMGs, as well as denotes metabolic genes that have been previously implicated in viral, non-host cell, metabolism (e.g. host cell entry) (**Supplementary File 4**).

Furthermore, DRAM-v flags metabolically active genes that baased on existing data are unlikely AMGs. This list includes CAZymes that are commonly used by viruses for host cell entry, peptidases that are commonly found in viruses according to MEROPs (5), viral structure and replication genes called by VOGDB, and if three metabolic genes are found in a single row (this is rare in viral genomes). Auxiliary scores for putative AMGs are based on the relative location on the vMAG of viral-like genes and viral hallmark genes, as determined by VirSorter (32). We recommend that users consider auxiliary scores 1-2 as high quality AMGs, category 3 as medium quality AMGs, and category 4-5 as low quality AMGs.

### SUPPLEMENTARY DISCUSSION

#### ***Comparison of DRAM features to other tools***

To highlight the value of DRAM, we have compared the features of DRAM to publically available annotators, genome veiwers, and metagenomic platforms (Supplementary Table 1).

I. DRAM is the only current annotation tool to provide an annotation framework for viruses, providing both a ruleset for defining auxiliary metabolic genes and an annotation system to functionally classfiy these putative genes.

II. DRAM is the only annotation tool to integrate taxonomic content and genome quality statistics. Moreover, this information is linked to the annotations, thus it is possible for DRAM users to assess quickly if a pathway is missing a key gene, if this is due to consistency with other closely related taxa, or due to genome incompleteness. Also, with the goal to improve genome quality standardization in the microbiome field, DRAM uses criteria established by Genome Standards Consortium (cite Bowers) and applies it to MAG lists, providing users with consistent interpretaion and content neeeded for genome deposition.

III. Beyond annotation, DRAM provides users with expertly curated metabolic summaries from multiple databases, see section "Interpreting results from DRAM and DRAM-v" above for details on the CAZyme and peptidases. While it may be possible for other annotation tools to utilize CAZyme and MEROPS databases, DRAM is the only tool that manually curates these databases, to provide users with putatitve substrate-use profiles for each genome.

IV. DRAM annotations enable comparative analyses of  thousands of genomes at once (specifically in the forms of the *Distillate* shown in Supplementary File 2 and the Product in Fig. 3). Other annotators such as METABOLIC provide curated metabolic summaries of genomes, but are more focused on single gene processes that link into biogeochemical cycles (e.g. nitrate reductase for nitrate reduction to nitrite), which DRAM also does but expands this fuction to provide more extensive pathway level completion for processes beyond biogeochemical cycles, such as flagellar assembly, nucleotide synthesis, antibiotic resistance, and transporters.

V. Custom marker inputs provided by the user can also be utilized in DRAM, as well as in some other platforms (e.g. Anvi'o, Metapathways, and METABOLIC).

VI. While Prokka and DFast only provide text file summaries, some tools, including DRAM do provide more comprehensive data visualization. For example, Metapathways and MetaErg visualizations allow users to visualize a single pathway in a single genome at one time, while METABOLIC visualization summaries are focused at the biogeochemical cycle level, quantifying the total sum of genomes (based on the presence of key functional genes) that may contribute to specific aspects of the sulfur, carbon, or nitrogen cycle. These visualizations while having strengths, are not effectively extended to hundreds or more genomes at once. DRAM is different from these tools and provides visual pathway summaries that can be used on thousands of genomes (*Product* Heatmap), allowing users to visually compare gene functional profiles for key biogeochemical reactions, as well as compare multiple pathways across multiple genomes at once.

It is important to note, that all of these annotation tools have inherent strengths and weaknesses that are based upon database selection and content, differences in output files, and in the case of some tools different data visualizations. Thus, it is likely that a robust annotation pipeline may simultaneoously use various tools in a complementary fashion.

***Measuring Precision and Recall of DRAM annotations using KEGG Genes and E. coli***

To further test the ability of DRAM to accurately annotate a bacterial genome we reannotated the *E. coli* K-12 MG1655 genome using filtered versions of the KEGG Genes database (2). While no gold standard set of annotations exist, we chose to use the *E. coli* K-12 MG1655 genome as it is one of the most completely biochemically characterized organisms. To evaluate the ability of DRAM to retrieve annotations the genome was annotated four times using 1) all DRAM databases including all of KEGG Genes (2), 2) the full set of DRAM databases with all genes from the *E. coli* K-12 MG1655 genomes removed from KEGG Genes (2), 3) the full set of DRAM databases with all genes from genomes in the

*E. coli* species removed from KEGG Genes (2) and 4) the full set of DRAM databases with all genes from genomes in the Escherichia genus removed from KEGG Genes (2). This allows us to test the ability of DRAM to recover annotations in genomes were close relatives are present and absent in the genome and evaluates whether DRAM is picking up false positive signals using default parameters (**Supplementary Methods**). DRAM showed high levels of performance across KEGG Genes (2) subsets (**Supplementary Figure 3**). Unsurprisingly performance was highest by all metrics when the genes from same genome were present but even when the entire genus of Escherichia was removed performance remained high with precision staying flat and recall falling as more genes were removed from the database. This suggests DRAM with default settings is relatively conservative and sacrifices recall for high levels of precision.

***Dechloromonas aromatica RCB full genome description***

To demonstrate the usefulness and data categorization of DRAM, we selected *Dechloromonas aromatica* RCB genome as an example (**Figure 2-3**). DRAM *Product* highlights that this genome has 7 of the 8 steps for glycolysis, corroborated in the *Distillate*, which shows that both types of phosphofructokinase are absent. Beyond summarizing genes per genome for pathway completion on the heatmap, the Distillate also displays genes involved in several other carbon and organic nitrogen metabolisms. In the *Decholoromonas aromatica* RCB genome, the *Distillate* (**Supplementary File 4**) shows the capacity for catechol meta-cleavage to acetyl-CoA, and urea degradation to ammonia. Collectively, DRAM has the capacity to annotate hundreds of MAGs at once, distilling genome annotations down to environmentally relevant metabolisms output to the DRAM-unique *Distillate* and *Product*.

***Integration of DRAM and DRAM-v to begin to infer virocell metabolism***

Integrated analyses using DRAM and DRAM-v offer a new opportunity to investigate the metabolic interplay that may exist between microbial hosts and viral symbionts. Here, to link metagenome derived hosts and viruses, we use CRISPR spacer matches between the HMP reconstructed MAGs and the HMP reconstructed vMAGs using (33). Four vMAGs had a perfect host match to a ~32 bp spacer sequence located in CRISPR-Cas arrays of four separate microbial MAGs. One of these four viruses and host linkages were of particular interest, as the viral genome encoded two adjacent medium-confidence AMGs and the host genome was an estimated 97% complete draft genome. This pairing offered a unique opportunity to investigate how virally encoded AMG metabolism may complement the host. The microbial host (bin.55) was a member of the genus *Lachnospira*, which are associated with pectin fermentation and short chain fatty acid production in mammalian guts (34, 35). Using DRAM, we inferred that this genome was an obligate fermenter and confirmed the presence of four pectin methylesterases (CE8) and two copies of three different pectate lyases (PL1, PL9, PL10), consistent with prior isolate characterizations (**Supplementary Figure 9**). Based on DRAM, we concluded that pectin degradation products could feed into a modified Entner-Doudoroff pathway via 2-dehydro-3-deoxy-phosphogluconate aldolase, generating reducing equivalents for fermentation to various end products. This *Lachnospira* genome also encodes a polyphenol oxidase that could enable it to depolymerize lignin, but appears to lack the capacity for subsequent downstream, monomeric lignin degradation.

We linked a 47,604 bp vMAG that was taxonomically identified as a mu-like virus in the family Myoviridae to this *Lachnospira* host genome. This vMAG encoded 62 genes, of which 80% were annotated as viral hallmark, viral-like, or virome exclusive, confirming the likelihood that this contig is of viral origin. The two putative AMGs on this vMAG were 4-oxalocrotonate tautomerase (4OT) and an alcohol dehydrogenase (ADH) and located adjacent to one another (**Supplementary Figure 9**). 4OT is involved in aerobic catechol meta-cleavage, a central aromatic compound catabolic pathway (36). However, as the host and the vMAG lack the key enzyme of the catechol meta-cleavage pathway (catechol 2,3-dioxygenase (37)), it is unlikely that the 4OT AMG is involved in this pathway. Recent studies have suggested that 4OT are promiscuous enzymes (36, 38), with detectable aldolase activity using aromatic aldehyde substrates (37). This is interesting for two reasons: promiscuity may increase the utility of the viral AMG, and this function may enable the host to ferment aromatic lignin monomers

(**Supplementary Figure 9**). The viral 4OT has 50-56% amino acid similarity to biochemically characterized 4OT enzymes, with three-dimensional protein sequence structure modelling giving high-confidence matches (>99.5%, **Supplementary Figure 10**) to various 4OT structures (**Supplementary File 4**). Sequence analysis reveals that the viral 4OT has key catalytic residues Pro1, Arg11, and Trp50, but Thr39 instead of Arg39 in homologs, however this residue is not involved in promiscuous aldolase catalysis. Therefore, we suggest that the viral 4OT may enable the host to carry out aldolase reactions with lignin monomers, producing acetaldehyde and aromatic aldehydes. These products can then be oxidized or reduced by the viral ADH, which has a three-dimensional protein sequence structural model that matches an aldehyde-alcohol dehydrogenase with high-confidence (100%, **Supplementary File 4**).

Summarizing, the DRAM-v identified AMGs, could complement *Lachnospira* metabolism, enabling the host to ferment previously inaccessible lignin monomers yielding aromatic aldehydes and alcohols, which could then be processed to generate alternative fermentation end products. While intriguing, we note that that future biochemical characterization of host and viral genes are needed to support this metabolic proposition. However, this example of a viral AMG lacking a homologue in the linked microbial host genome opens new research directions of how viruses may expand the metabolic repertoires of their hosts, with implications for host-virus coevolution and reshaping of the human microbial gut chemical landscape.

Protein sequences from the two putative host-linked AMGs (ADH: HMP1_viralSeqs_398_VIRSorter_scaffold_938-cat_2_58 and 4OT: HMP1_viralSeqs_398_VIRSorter_scaffold_938-cat_2_59) were structurally modeled using PHYRE2 (38) to validate DRAM-v functional predictions. There were 100% and 99.7% confidence scores to bacterial enzymes matching alcohol dehydrogenase and 4-oxalocrotonate tautomerase. Both AMGs were blasted to UniProt (19) reviewed protein entries and had best-matches to biochemically studied alcohol dehydrogenase and 2-hydroxymuconate tautomerase (synonym 4-oxalocrotonate tautomerase). Catalytic residues in 4OT were manually curated. See **Supplementary File 4** for Phyre2 (39) and UniProt (19) hits.

**SUPPLEMENTARY TABLES:**

| ***Annotator*** | *Curated metabolic summaries* | *Custom marker inputs* | *Annotation visuali-zations* | *Annotates with multiple databases* | *CAZyme curation and synthesis* | *Peptidase curation and synthesis* | *Scales beyond 500 genomes* | *Provides genome quality summarys* | *Viral genome analysis-AMGs* |
|---|---|---|---|---|---|---|---|---|---|
| ***DRAM*** | **X** | **X** | **X** | **X** | **X** | **X** | **X** | **X** | **X** |
| *Prokka* (9) | | | | X | | | X | | |
| *DFAST* (10) | | | | X | | | X | | |
| *MetaERG* (11) | | | X | X | | | X | | |
| *Anvi'o* (40) | | X | | X | | | | | |
| *GenomeProperties* (41) | X | | X | | | | | | |
| *Metapathways* (42) | | X | X | X | | | X | | |
| *METABOLIC* (43) | X | X | X | X | | | X | | |

**Supplementary Table 1: Comparison of features across several annotators, with these topics discusssed more in section "Comparison of DRAM features" above.**

### SUPPLEMENTARY FIGURES:

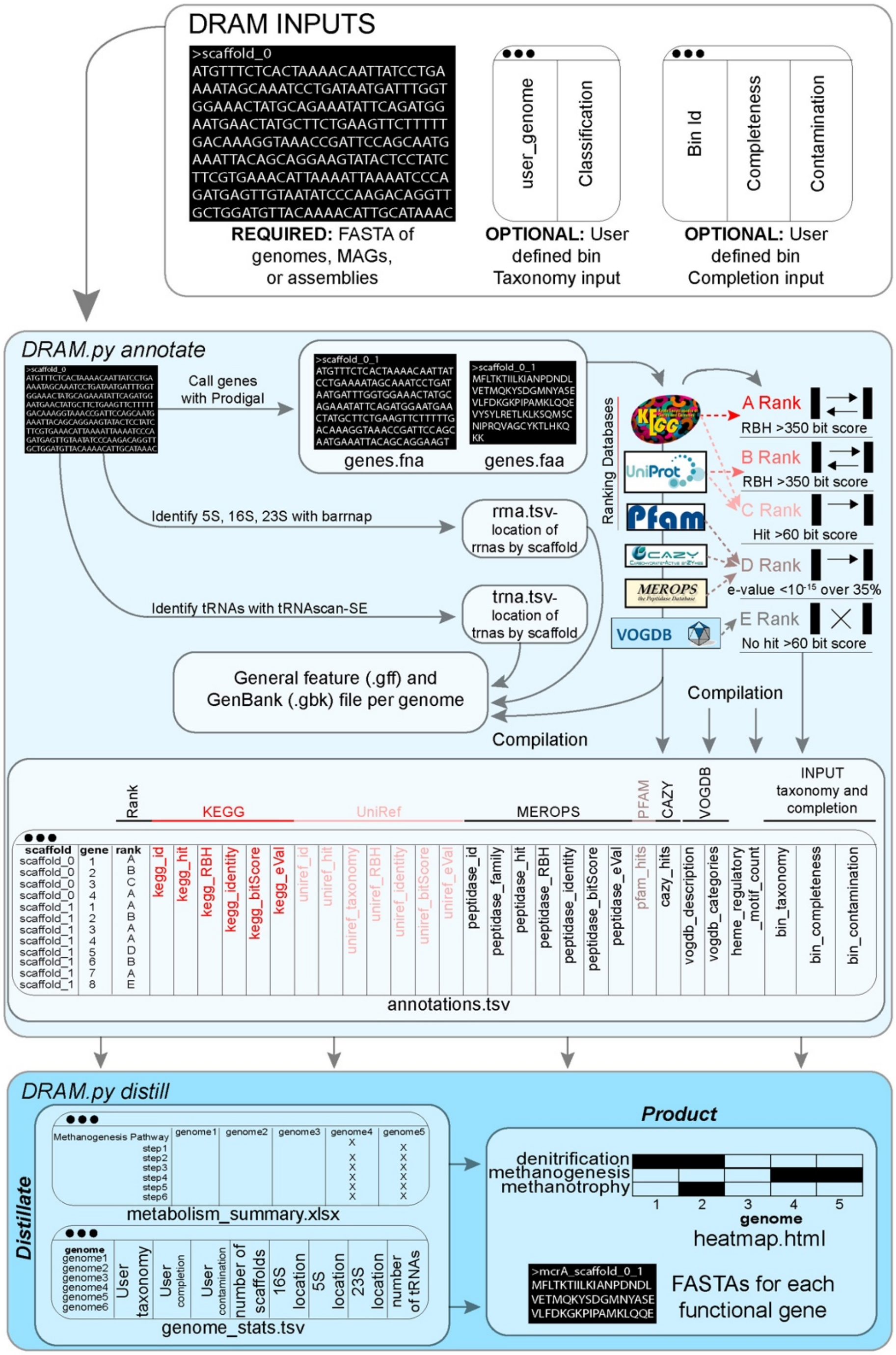


**Supplementary Figure 1: Detailed overview of DRAM pipeline.** Workflow highlights DRAM inputs and outputs, as well as how all ranks, annotations, and analyses are compiled together.

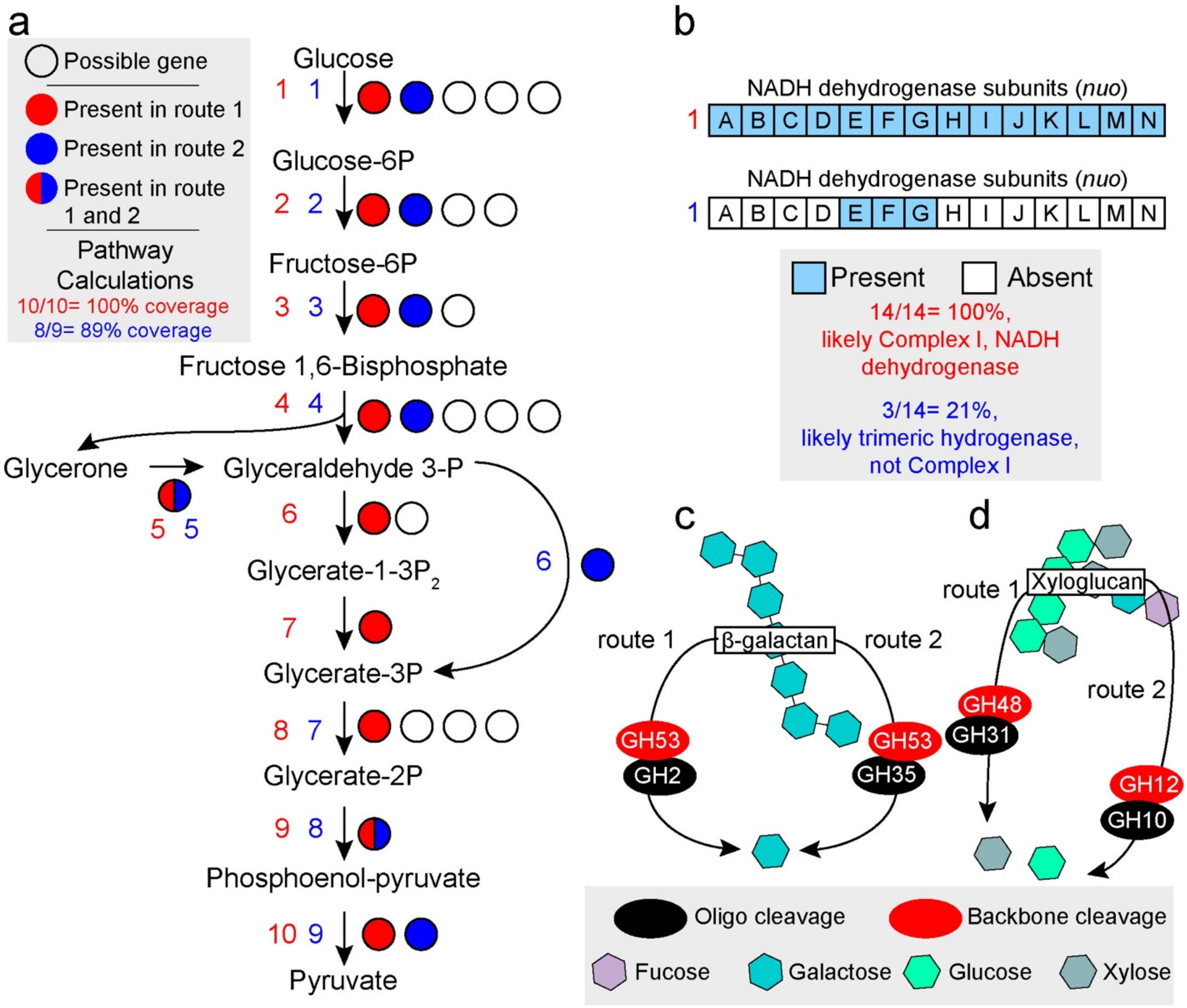


**Supplementary Figure 2: Pathway completion and redundancy/exclusivity logic used in DRAM. a** Pathway completion considers alternate routes for processes that start and end with the same compounds, while also accounting for different genes (shown by black circle outlines) at each step. In this case, glycolysis can occur via a 10 (red 1-10) or 9 (blue 1-9) step pathway, with each step having alternate genes possible. Pathway completion is calculated based on each route individually. Shown here, the red pathway has 10 out of 10 steps, so completion would be reported as 100%. The blue pathway would be reported as 89% (8 out of 9 steps) because it is missing step 7. **b** The value of our enzyme complex completion for reducing misannotations is demonstrated here, as the partial completion (3 genes) of the multi-subunit NADH dehydrogenase is not due to a complete complex I, but rather the presence of hydrogenases common in obligate fermenters. These hydrogenases are further annotated in detail by their type and function in *Distillate* (**Supplementary File 2**). **c** Some GHs are redundant, meaning they degrade the same substrate, thus an organism needs to only have one type to degrade a particular carbohydrate. Shown here, to degrade B-galactan, organisms can have GH53 and GH2 (route 1) OR GH53 and GH35 (route 2). **d** In this example, xyloglucan requires GH48 and GH31 OR GH12 and GH10.

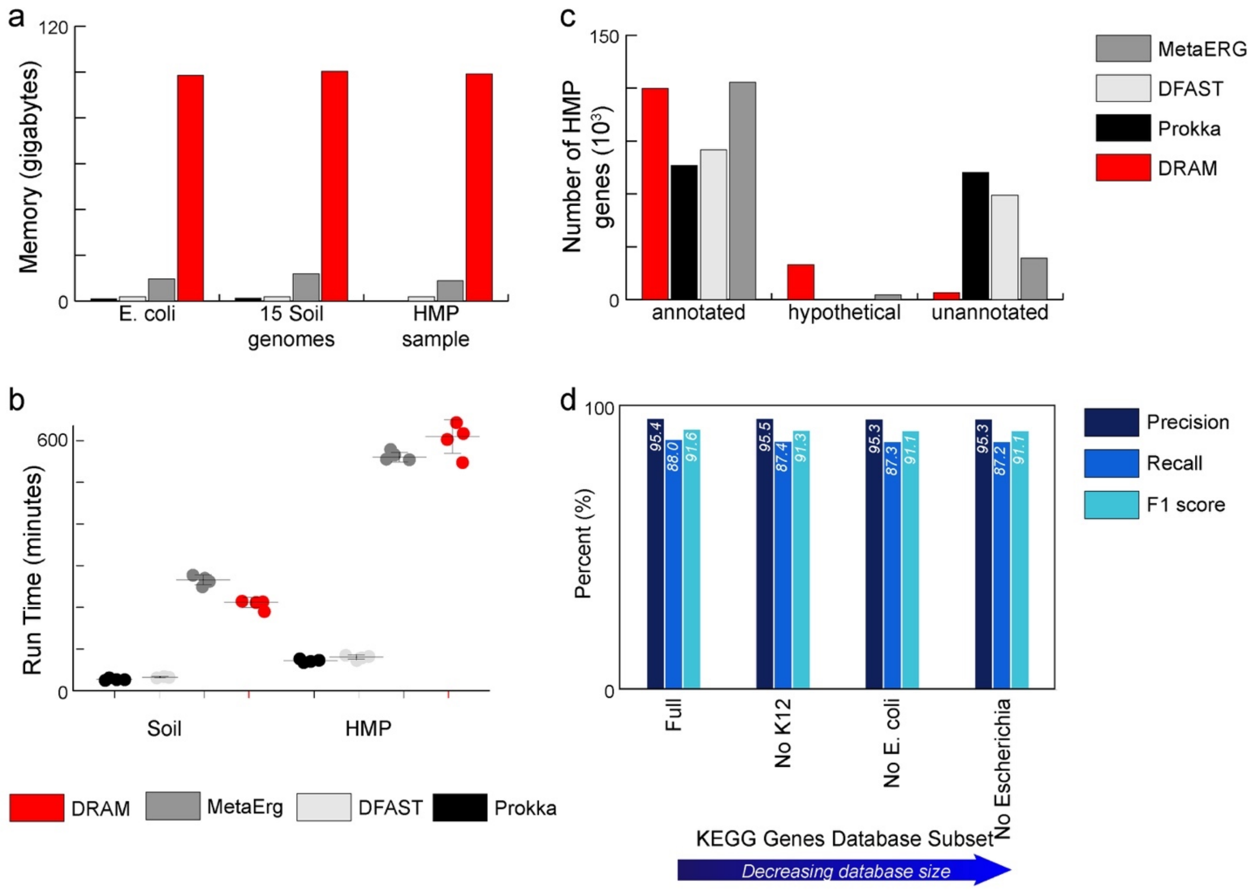


**Supplementary Figure 3: DRAM performance. a.** Barcharts showing maximum memory usage for Prokka (9), DFAST (10), MetaErg (11), and DRAM across three different files. All annotations were run with 10 processors. **b.** Dot plot shows the run time of 4 replicate runs for each annotator on both *in silico* soil and HMP genomes, with horizontal lines noting the median and error bars noting one quartile from the median. **c.** Barcharts in number of annotated, hypothetical, and unannotated genes assigned by each annotator when analyzing HMP gut metagenome derived MAGs. See methods for definitions of annotated, hypothetical, and unannotated genes, relative to each annotator. **d.** Bar charts showing the precision, recall and F1 score based on recovery of KEGG Orthology annotation from the *E. coli* K12 MG1655 genome using DRAM with different subsets of the KEGG Genes database removed. The full bars represent performance when the full KEGG Genes database (2) was used. No K12 represents performance when the genes from E. coli K12 MG1655 were removed from the KEGG Genes database. No E. coli shows performance when all genes from all genomes of the *E. coli* species were removed. No

*Escherichia* represents performance when all genes from all genomes from all members of the *Escherichia* genus were removed.

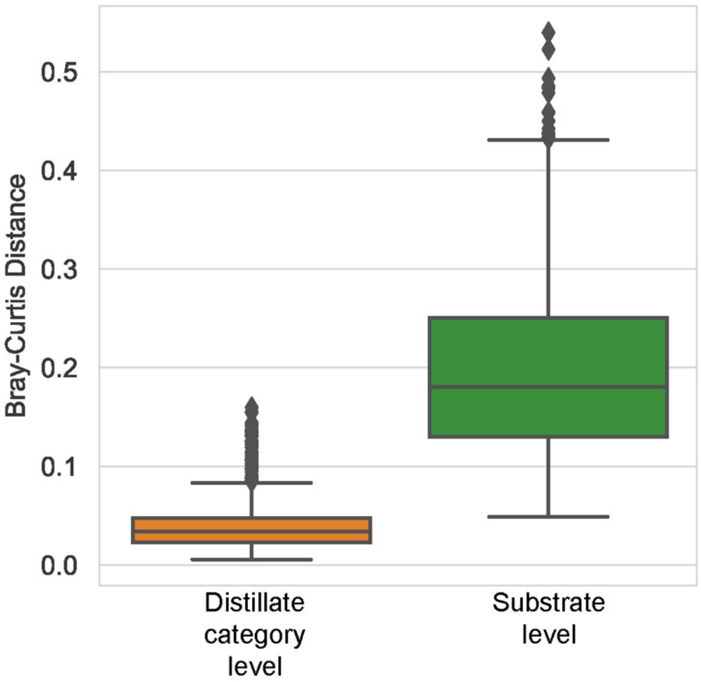


**Supplementary Figure 4: Beta diversity measures comparing distillate category diversity to carbohydrate diversity.** The beta diversity as measured by Bray-Curtis distance between all pairs of 44 HMP metagenomes at the DRAM header (*Distillat*e) and substrate utilization (Substrate) level, demonstrating that substrate level variability across humans is significantly higher than at bulk level annotation categories (*Distillate*) ($p<1x10^{-25}$).

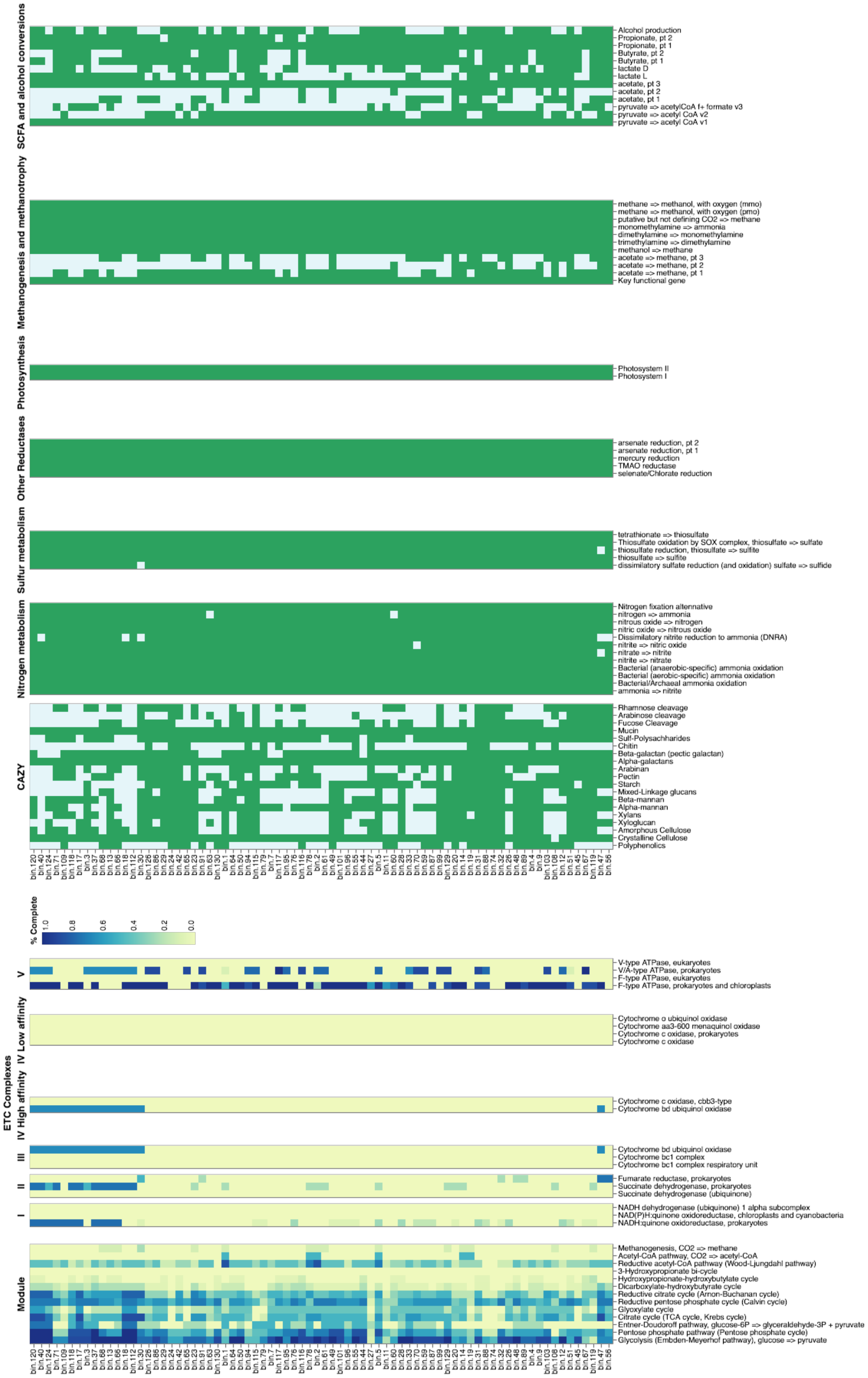


**Supplementary Figure 5: Full *Product* heatmap for all Human Microbiome Project from Human Microbiome Project (HMP) reconstructed genomes Figure 5.** Heatmap shows the DRAM *Product* output for 76 HMP MAGs reconstructed in this study.

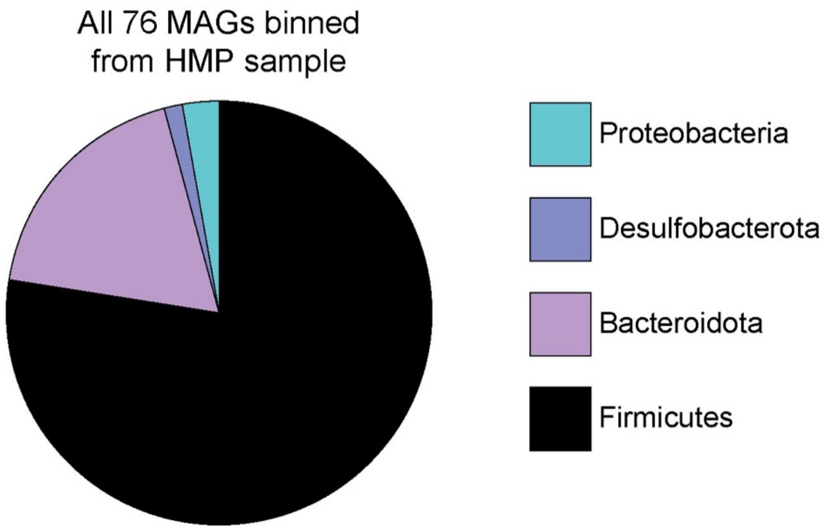


**Supplementary Figure 7: Reconstructed genomes from Human Microbiome Project metagenomes reflect the composition typically recovered in the human gut.** Piechart shows the percent of bins by Phylum level taxonomy.

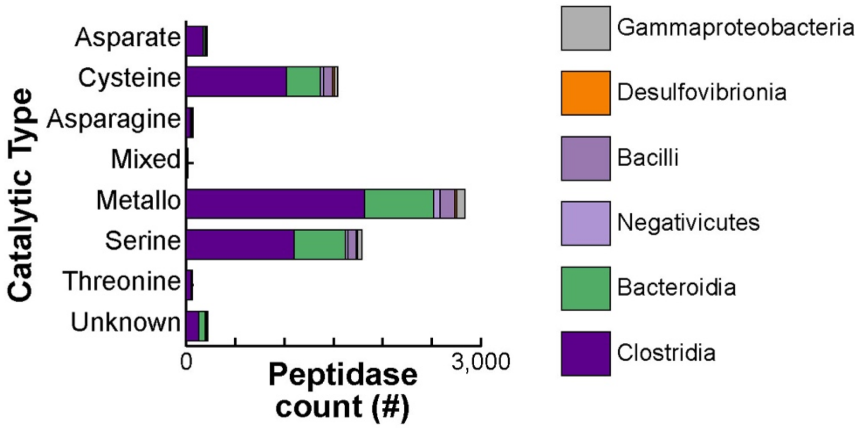


**Supplementary Figure 8: Peptidases are prevalent across 76 genomes from Human Microbiome Project (HMP) reconstructed genomes.** Stacked bar chart of peptidases recovered from 76 HMP MAGs reported in Figure 5. Bars represent peptidase catalytic type, summed from the MEROPs (5) database. The bars are colored by genome taxonomy class in which the gene was derived.

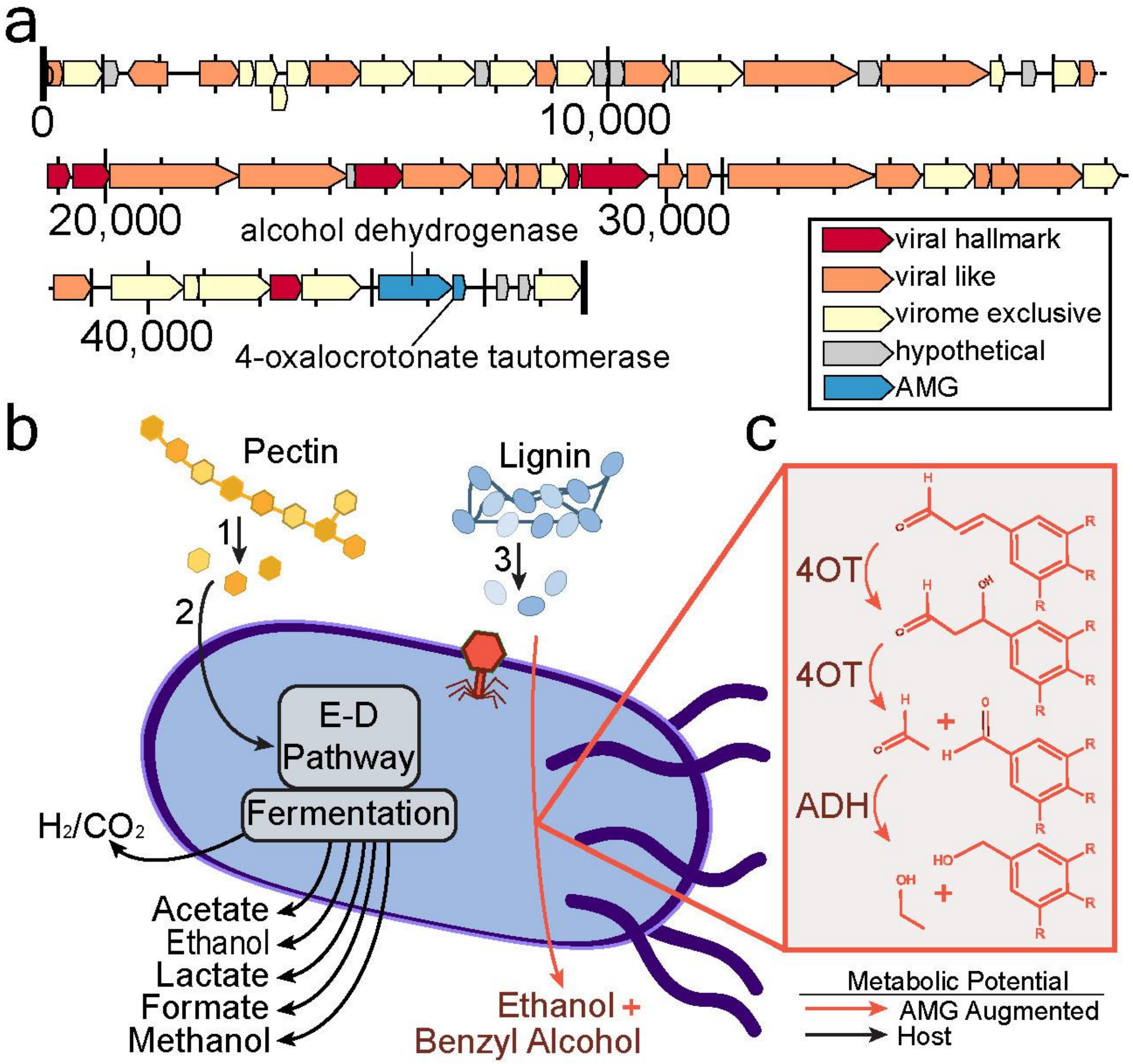


**Supplementary Figure 9: Integrated use of DRAM and DRAM-v to develop hypotheses about virocell metabolism. a** VMAG recovered from HMP with CRISPR links to a MAG encodes two putative AMGs for carbon metabolism. Open Reading Frames (ORFs) are represented by arrows and are colored by viral function. **b** DRAM-identified *Lachnospira*-encoded potential for pectin fermentation and lignin depolymerization are represented by black arrows. Numbers correspond to encoded enzymes as follows: (1) PL1, PL9, PL10, CE8, CE12; (2) GH105, PL26, GH28, (3) polyphenol oxidase-laccase. **f** DRAM-v identified putative auxiliary metabolic genes transformations encoded by a *Lachnospira* CRISPR-linked vMAG are shown in red. Briefly, 4-oxalocrotonate tautomerase (4OT) is inferred to carry out subsequent aldolase reactions on lignin-derived aromatic aldehydes, generating acetaldehyde and benzaldehydes. The viral encoded alcohol-aldehyde dehydrogenase (ADH) then can reduce (as shown) or oxidize these products.

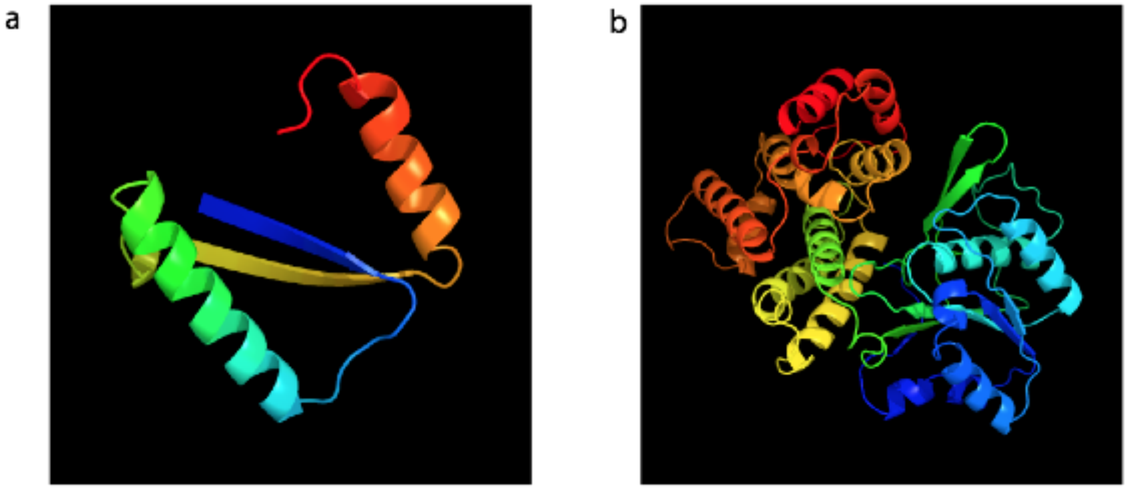


**Supplementary Figure 10**: **Phyre2 (39) structural models for *Lachnospira*-linked AMGs. a** 4-oxalocrotonate tautomerase and **b** alcohol dehydrogenase. N-terminal residues are red, and C-terminal residues are blue. Phyre2 (39) structural model information in **Supplementary file 4**.

**SUPPLEMENTARY FILES:**

**Supplementary File 1:** DRAM *Raw* annotation output from HMP MAG annotations, with each row representing a called gene and each column noting specific annotation information about each gene. The first seven columns of this file contain information on gene calling of input scaffolds or contigs, with the first column summarizing each called gene with a unique identifier—fastainputfilename_scaffoldname_calledgenenumber. The subsequent columns (2-4), break this information down into FASTA input file name, scaffold name, and gene position (number). Columns two 5-7 denote the base pair location start and end position for each gene on the scaffold as well as the strandedness from Prodigal. Annotation information of each called gene is stored in columns 8-24. Specifically, KEGG outputs (KEGG id (KO number), KEGG annotation, Reverse Best Hit (RBH) reported as True or False, percent identity match, bitscore, e-value) for each gene are stored in columns 8-13, MEROPS outputs (MEROPS id number, MEROPS peptidase family, MEROPS annotation, Reverse Best Hit (RBH) reported as True or False, percent identity match, bitscore, e-value) for each gene are stored in columns 14-20, best Pfam hits are store in column 21, best CAZY hits in column 22, and best VOGDB hits and category in columns 23-24. Column 25 contains the number of CXXCH motifs per gene to expedite classifying multi-heme cytochromes. Lastly, if a user provided taxonomy and quality information for each input FASTA file, these are stored in columns 26-28.

**Supplementary File 2:** DRAM *Distillate* output from HMP MAG annotations, with each tab representing a metabolism category, rows presenting specific genes within each metabolism category, and columns representing each input FASTA (in this case each MAG).

**Supplementary File 3:** Full DRAM *Product* output in html format from soil isolate and MAG annotations shown in main text Figure 3.

**Supplementary File 4:** Excel tables for:

(Tab 1): Database size comparisons of DRAM, Prokka (9), DFAST (10), and MetaErg (11).

(Tab 2): Performance comparisons of DRAM, Prokka (9), DFAST (10), and MetaErg (11) per dataset.

(Tab 3): Performance comparisons of Prokka (9), DFAST (10), and MetaErg (11) per soil genome.

(Tab 4): Soil isolate and MAG statistics according to Bowers, et al. (6), including accession numbers.

(Tab 5): Raw data for E. coli KEGG gene dropout tests used to demonstrate accuracy of annotations in DRAM, with data visualized in **Supplementary Figure 3**.

(Tab 6): HMP assembly stats and accession numbers.

(Tab 7): HMP MAG statistics according to Bowers, et al (6).

(Tab 8): Viral genome statistics according to Roux et al (8).

(Tab 9): Viral AMG database, including the details on 17 experimentally verified AMGs.

(Tab 10): *Product* logic including CAZyme calls.

(Tab 11): *Distillate* logic including non-AMG calls.

(Tab 12): Gene annotations corresponding to *Dechloromonas aromatica* RCB metabolic map in **Figure 2a**.

(Tab 13): Phyre2 (39) structural model information for AMG in **Supplementary Figure 9-10**.

**Supplementary File 5:** DRAM *Distillate* output from soil isolate genomes and MAGs.

**Supplementary File 6:** DRAM-v *Distillate* output from soil vMAGs from Emerson, et al (15).

**Supplementary File 7:** DRAM-v *Product* output from soil vMAGs from Emerson, et al (15).

**Supplementary File 8:** DRAM-v *Distillate* output from HMP derived vMAGs.

**Supplementary File 9:** DRAM-v *Product* output from HMP derived vMAGs.

**Supplementary File 10:** FASTA file of viral genomes from Emerson, et al. (15) and recovered here from HMP metagenomes.

### SUPPLEMENTARY REFERENCES

1. El-Gebali,S., Mistry,J., Bateman,A., Eddy,S.R., Luciani,A., Potter,S.C., Qureshi,M., Richardson,L.J., Salazar,G.A., Smart,A., *et al.* (2019) The Pfam protein families database in 2019. *Nucleic Acids Res.*, **47**, D427–D432.

2. Kanehisa,M., Furumichi,M., Tanabe,M., Sato,Y. and Morishima,K. (2017) KEGG: new perspectives on genomes, pathways, diseases and drugs. *Nucleic Acids Res.*, **45**, D353–D361.

3. Suzek,B.E., Wang,Y., Huang,H., McGarvey,P.B. and Wu,C.H. (2015) UniRef clusters: a comprehensive and scalable alternative for improving sequence similarity searches. *Bioinformatics*, **31**, 926–932.

4. Zhang,H., Yohe,T., Huang,L., Entwistle,S., Wu,P., Yang,Z., Busk,P.K., Xu,Y. and Yin,Y. (2018) DbCAN2: A meta server for automated carbohydrate-active enzyme annotation. *Nucleic Acids Res.*, **46**, W95–W101.

5. Rawlings,N.D., Barrett,A.J., Thomas,P.D., Huang,X., Bateman,A. and Finn,R.D. (2018) The MEROPS database of proteolytic enzymes, their substrates and inhibitors in 2017 and a comparison with peptidases in the PANTHER database. *Nucleic Acids Res.*, **46**, D624–D632.

6. Bowers,R.M., Kyrpides,N.C., Stepanauskas,R., Harmon-Smith,M., Doud,D., Reddy,T.B.K., Schulz,F., Jarett,J., Rivers,A.R., Eloe-Fadrosh,E.A., *et al.* (2017) Minimum information about a single amplified genome (MISAG) and a metagenome-assembled genome (MIMAG) of bacteria and archaea. *Nat. Biotechnol.*, **35**, 725–731.

7. Pruitt,K.D., Tatusova,T. and Maglott,D.R. (2007) NCBI reference sequences (RefSeq): a curated non-redundant sequence database of genomes, transcripts and proteins. *Nucleic Acids Res.*, **35**, D61--D65.

8. Roux,S., Adriaenssens,E.M., Dutilh,B.E., Koonin,E. V., Kropinski,A.M., Krupovic,M., Kuhn,J.H., Lavigne,R., Brister,J.R., Varsani,A., *et al.* (2019) Minimum information about an uncultivated virus genome (MIUVIG). *Nat. Biotechnol.*, **37**, 29–37.

9. Seemann,T. (2014) Prokka: rapid prokaryotic genome annotation. *Bioinformatics*, **30**, 2068–2069.

10. Tanizawa,Y., Fujisawa,T. and Nakamura,Y. (2018) DFAST: a flexible prokaryotic genome annotation pipeline for faster genome publication. *Bioinformatics*, **34**, 1037–1039.

11. Dong,X. and Strous,M. (2019) An Integrated Pipeline for Annotation and Visualization of Metagenomic Contigs. *Front. Genet.*, **10**, 999.

12. Gevers,D., Knight,R., Petrosino,J.F., Huang,K., McGuire,A.L., Birren,B.W., Nelson,K.E., White,O., Methé,B.A. and Huttenhower,C. (2012) The Human Microbiome Project: a community resource for the healthy human microbiome. *PLoS Biol.*, **10**, e1001377.

13. Huttenhower,C., Gevers,D., Knight,R., Abubucker,S., Badger,J.H., Chinwalla,A.T., Creasy,H.H., Earl,A.M., Fitzgerald,M.G., Fulton,R.S., *et al.* (2012) Structure, function and diversity of the healthy human microbiome. *Nature*, **486**, 207–214.

14. Peng,Y., Leung,H.C.M., Yiu,S.M. and Chin,F.Y.L. (2012) IDBA-UD: a de novo assembler for single-cell and metagenomic sequencing data with highly uneven depth. *Bioinformatics*, **28**, 1420–1428.

15. Emerson,J.B., Roux,S., Brum,J.R., Bolduc,B., Woodcroft,B.J., Jang,H. Bin, Singleton,C.M., Solden,L.M., Naas,A.E., Boyd,J.A., *et al.* (2018) Host-linked soil viral ecology along a permafrost thaw gradient. *Nat. Microbiol.*, **3**, 870–880.

16. Anantharaman,K., Brown,C.T., Hug,L.A., Sharon,I., Castelle,C.J., Probst,A.J., Thomas,B.C., Singh,A., Wilkins,M.J., Karaoz,U., *et al.* (2016) Thousands of microbial genomes shed light on interconnected biogeochemical processes in an aquifer system. *Nat. Commun.*, **7**, 13219.

17. Bairoch,A. and Boeckmann,B. (1991) The SWISS-PROT protein sequence data bank. *Nucleic Acids Res.*, **19**, 2247.

18. Haft,D.H., Selengut,J.D. and White,O. (2003) The TIGRFAMs database of protein families. *Nucleic Acids Res.*, **31**, 371–373.

19. UniProt: the universal protein knowledgebase (2017) *Nucleic Acids Res.*, **45**, D158--D169.

20. Chaumeil,P.-A., Mussig,A.J., Hugenholtz,P. and Parks,D.H. (2018) GTDB-Tk: a toolkit to classify genomes with the Genome Taxonomy Database. *Bioinformatics*, **36**, 1925–1927.

21. Wrighton,K.C., Thomas,B.C., Sharon,I., Miller,C.S., Castelle,C.J., VerBerkmoes,N.C., Wilkins,M.J.,

Hettich,R.L., Lipton,M.S., Williams,K.H., *et al.* (2012) Fermentation, hydrogen, and sulfur metabolism in multiple uncultivated bacterial phyla. *Science (80-. ).*, **337**, 1661–1665.

22. Vignais,P.M. and Billoud,B. (2007) Occurrence, classification, and biological function of hydrogenases: An overview. *Chem. Rev.*, **107**, 4206–4272.

23. Parks,D.H., Imelfort,M., Skennerton,C.T., Hugenholtz,P. and Tyson,G.W. (2015) CheckM: assessing the quality of microbial genomes recovered from isolates, single cells, and metagenomes. *Genome Res.*, **25**, 1043–1055.

24. Chan,P.P. and Lowe,T.M. (2019) tRNAscan-SE: Searching for tRNA genes in genomic sequences. In *Methods in Molecular Biology*. Humana Press Inc., Vol. 1962, pp. 1–14.

25. Yu,N.Y., Wagner,J.R., Laird,M.R., Melli,G., Rey,S., Lo,R., Dao,P., Sahinalp,S.C., Ester,M., Foster,L.J., *et al.* (2010) PSORTb 3.0: improved protein subcellular localization prediction with refined localization subcategories and predictive capabilities for all prokaryotes. *Bioinformatics*, **26**, 1608–1615.

26. Danczak,R.E., Johnston,M.D., Kenah,C., Slattery,M., Wrighton,K.C. and Wilkins,M.J. (2017) Members of the Candidate Phyla Radiation are functionally differentiated by carbon-and nitrogen-cycling capabilities. *Microbiome*, **5**, 112.

27. Ticak,T., Kountz,D.J., Girosky,K.E., Krzycki,J.A. and Ferguson,D.J. (2014) A nonpyrrolysine member of the widely distributed trimethylamine methyltransferase family is a glycine betaine methyltransferase. *Proc. Natl. Acad. Sci.*, **111**, E4668--E4676.

28. Srinivasan,G., James,C.M. and Krzycki,J.A. (2002) Pyrrolysine encoded by UAG in Archaea: charging of a UAG-decoding specialized tRNA. *Science (80-. ).*, **296**, 1459–1462.

29. Solden,L.M., Naas,A.E., Roux,S., Daly,R.A., Collins,W.B., Nicora,C.D., Purvine,S.O., Hoyt,D.W., Schückel,J., Jørgensen,B., *et al.* (2018) Interspecies cross-feeding orchestrates carbon degradation in the rumen ecosystem. *Nat. Microbiol.*, **3**, 1274.

30. Koropatkin,N.M., Cameron,E.A. and Martens,E.C. (2012) How glycan metabolism shapes the human gut microbiota. *Nat. Rev. Microbiol.*, **10**, 323–335.

31. Grondin,J.M., Tamura,K., Déjean,G., Abbott,D.W. and Brumer,H. (2017) Polysaccharide utilization loci: fueling microbial communities. *J. Bacteriol.*, **199**, e00860--16.

32. Roux,S., Enault,F., Hurwitz,B.L. and Sullivan,M.B. (2015) VirSorter: mining viral signal from microbial genomic data. *PeerJ*, **3**, e985.

33. Daly,R.A., Roux,S., Borton,M.A., Morgan,D.M., Johnston,M.D., Booker,A.E., Hoyt,D.W., Meulia,T., Wolfe,R.A., Hanson,A.J., *et al.* (2019) Viruses control dominant bacteria colonizing the terrestrial deep biosphere after hydraulic fracturing. *Nat. Microbiol.*, **4**, 352–361.

34. Bang,S.-J., Kim,G., Lim,M.Y., Song,E.-J., Jung,D.-H., Kum,J.-S., Nam,Y.-D., Park,C.-S. and Seo,D.-H. (2018) The influence of in vitro pectin fermentation on the human fecal microbiome. *Amb Express*, **8**, 1–9.

35. Dušková,D. and Marounek,M. (2001) Fermentation of pectin and glucose, and activity of pectin-degrading enzymes in the rumen bacterium Lachnospira multiparus. *Lett. Appl. Microbiol.*, **33**, 159–163.

36. Burks,E.A., Fleming,C.D., Mesecar,A.D., Whitman,C.P. and Pegan,S.D. (2010) Kinetic and structural characterization of a heterohexamer 4-oxalocrotonate tautomerase from chloroflexus aurantiacus J-10-fl: Implications for functional and structural diversity in the tautomerase superfamily. *Biochemistry*, **49**, 5016–5027.

37. Harayama,S. and Rekik,M. (1989) Bacterial aromatic ring-cleavage enzymes are classified into two different gene families. *J. Biol. Chem.*, **264**, 15328–33.

38. Davidson,R., Baas,B.J., Akiva,E., Holliday,G.L., Polacco,B.J., LeVieux,J.A., Pullara,C.R., Zhang,Y.J., Whitman,C.P. and Babbitt,P.C. (2018) A global view of structure–function relationships in the tautomerase superfamily. *J. Biol. Chem.*, **293**, 2342–2357.

39. Kelley,L.A., Mezulis,S., Yates,C.M., Wass,M.N. and Sternberg,M.J.E. (2015) The Phyre2 web portal for protein modeling, prediction and analysis. *Nat. Protoc.*, **10**, 845–858.

40. Eren,A.M., Esen,Ö.C., Quince,C., Vineis,J.H., Morrison,H.G., Sogin,M.L. and Delmont,T.O. (2015) Anvi'o: an advanced analysis and visualization platform for 'omics data. *PeerJ*, **3**, e1319.

41. Richardson,L.J., Rawlings,N.D., Salazar,G.A., Almeida,A., Haft,D.R., Ducq,G., Sutton,G.G. and Finn,R.D. (2019) Genome properties in 2019: a new companion database to InterPro for the inference of complete functional attributes. *Nucleic Acids Res.*, **47**, D564--D572.

42. Konwar,K.M., Hanson,N.W., Pagé,A.P. and Hallam,S.J. (2013) MetaPathways: a modular pipeline for constructing pathway/genome databases from environmental sequence information. *BMC Bioinformatics*, **14**, 202.

43. Zhou,Z., Tran,P., Liu,Y., Kieft,K. and Anantharaman,K. (2019) METABOLIC: A scalable high-throughput metabolic and biogeochemical functional trait profiler based on microbial genomes. *bioRxiv*.
